# Comprehensively Testing the Function of Missense Variation in the *STK11* Tumour Suppressor

**DOI:** 10.1101/2025.07.14.664734

**Authors:** Daniel Zimmerman, Atina Cote, Warren van Loggerenberg, Marinella Gebbia, Nishka Kishore, Jochen Weile, Roujia Li, Chloe Reno, Ashley PL Marsh, Felicia Hernandez, Preksha Shahagadkar, Lauren Grove, Samuel R. Meier, Hsin-Jung Wu, Silvia Fenoglio, Leanne Ahronian, Teng Teng, Andrew J Waters, David Seward, Mikko Taipale, Melyssa Aronson, Marcy E Richardson, David J Adams, Frederick P Roth

## Abstract

The tumor suppressor gene *STK11* encoding Serine/Threonine Kinase 11 (STK11) is associated with Peutz-Jeghers Syndrome (PJS), a heritable gastrointestinal disease that increases lifetime cancer risk, and with somatic variation that contributes to ∼30% of lung and 20% of cervical cancers. Although identifying pathogenic variants is clinically actionable, over 94% of *STK11* missense variants that have been observed clinically lack a definitive classification. We therefore measured the impact of *STK11* variants at scale in a mammalian cell-based assay, scoring 6,026 (73% of all possible) amino acid substitutions across the full-length gene. Functional scores—which were consistent with biochemical properties, smaller-scale assays, and pathogenicity annotations—identified a subset of PJS patients with germline *STK11* variants diagnosed later in life, as well as somatic *STK11* variants found in cancer patients that had comparable overall survival estimates to wild-type *STK11*. Our scores provided new evidence for 350 annotated VUS *STK11* missense variants and ∼80% of missense variants that have not yet been reported clinically, but we might expect to observe in the future. Thus, our effect map provides a proactive resource for gaining sequence-structure-function insights and evidence for actionable interpretation of clinical missense variants.

## Main

The serine/threonine kinase STK11 (also known as LKB1) phosphorylates multiple protein targets, including AMPK, which in turn inhibits the mTOR complex and reduces cell proliferation^1^. STK11 serves as a tumour suppressor in lung, cervical, and many other cancers. For example, somatic loss-of-function variants in *STK11* are found in >20% of non-small cell lung cancer (NSCLC) patients, where they are associated with poorer prognosis and resistance to immunotherapy^2,3^. Germline loss-of-function variants are associated with the rare (incidence of 1 in 50-200,000)^4^ monogenic disease PJS, which confers an increased lifetime risk of cancer in multiple tissue types (∼21% of individuals with PJS develop cancer by age 40 and ∼71% by age 70)^5^, and is characterized by growth of hamartomatous polyps in the small and large intestine^6^. Germline genetic testing offers a non-invasive means for early PJS diagnosis and improved clinical outcomes through heightened surveillance (e.g. colonoscopy).

Unfortunately, of 1,112 (ClinVar reported)^7^ missense variants, ∼94% (1,050) have been reported as either a variant of uncertain significance (VUS) (910) or have conflicting classifications (140) (Fig.1a), limiting the potential for improved care based on genetic testing. Similarly, limited knowledge of the pathogenicity of missense variants in *STK11* may cause a subset of NSCLC patients to miss potentially helpful therapies that are in development for *STK11*-mutated cancers.

Because functional assays can provide strong evidence to more definitively classify clinical missense variants^8–10^, and because every possible single-nucleotide change in *STK11* is likely to already exist in the human population^11^, systematically assessing missense variant effects on *STK11* function could serve to resolve the pathogenicity of many human variants (in a manner agnostic to ethnicity), including those that have yet to be clinically observed.

Here we set out to characterize all possible *STK11* variants via a multiplexed proliferation assay in human cells, generating the largest and most comprehensive resource on functional variation in *STK11* to date. Results were validated against biochemical properties, alternative functional assays (including *STK11* essentiality in HAP1 cells and enrichment in mouse-xenografted A549 tumours), known pathogenic and benign variants, and clinical phenotypes. Functional scores were calibrated using a Bayesian approach to derive evidence strength measures for use within the ACMG variant interpretation framework, thus providing new evidence towards pathogenicity or benignity for 5,863 STK11 amino acid substitutions (including 68% of clinically reported variants that are currently classified as VUS).

## Results

### A scalable assay for STK11 function

Given the scalability and accuracy of cell proliferation-based assays of variant impact that have been demonstrated for other proteins^12,13^, we sought such an assay for STK11. We built on previous observations that HeLa cells do not express functional STK11 (due to an Alu-element mediated homozygous deletion of the first three exons^14^), that introduction of a functional wild-type copy of *STK11* into HeLa cells reduces their growth rate^15–17^, and that HeLa growth upon expression of different *STK11* variants can distinguish normally functioning *STK11* variants from those that are damaging and disruptive of gene function^18^.

To both reproduce such an assay and demonstrate that it can be multiplexed, we generated a HeLa derived cell line bearing a Bxb1 ‘landing pad’^19^ at a safe harbor site, identified a small set of reference variants comprised of p.K78I, a known kinase dead variant, p.W239C and p.W308C, variants expected to impact protein-protein interactions, p.L286*, an early truncation, and expected tolerated variants p.I322L and p.S404F^20,21^. We generated the mutant *STK11* constructs in vectors suitable for integration via Bxb1 recombinase, and transfected these together (see Methods). The results showed clear separation of known pathogenic and benign variants after two weeks (Supplementary Fig. 1), giving us confidence to move forward with this assay at a larger scale.

### Generahting a missense variant effect map for STK11

To characterize nearly all possible *STK11* variants, we carried out a multiplexed proliferation assay in human cells using the following deep mutational scanning workflow: First, we generated an *STK11* variant library, employing a pool of oligonucleotides, each bearing a central three-nucleotide-position degeneracy corresponding to a given codon within the *STK11* cDNA, such that each clone contains approximately one position with a random codon replacement. Second, we shuttled the *STK11* variant library into a vector for integration in HeLa cells using a large-scale recombinational cloning procedure.^22^ Third, we generated a pool of cells each carrying a single *STK11* cDNA variant at a genomic landing pad in HeLa cells, using large-scale pooled transfection, recombinase-driven plasmid integration, and enrichment for integrant cells via fluorescence-activated cell sorting (FACS). Fourth, we performed growth-based selection in standard nutrient-rich media. Fifth and finally, we quantified the frequencies of each variant, both before and after growth selection, using next-generation sequencing (NGS) of >2M reads from each of 12 ∼110nt ‘tiles’ that collectively spanned the *STK11* cDNA (Fig. 1b). We performed three independent biological replicates (i.e. separate transfections of the *STK11* variant libraries) of the *STK11*-HeLa proliferation assay, each with two technical replicates (i.e. separate lineages of the same initial transfection).

**Figure 1.**
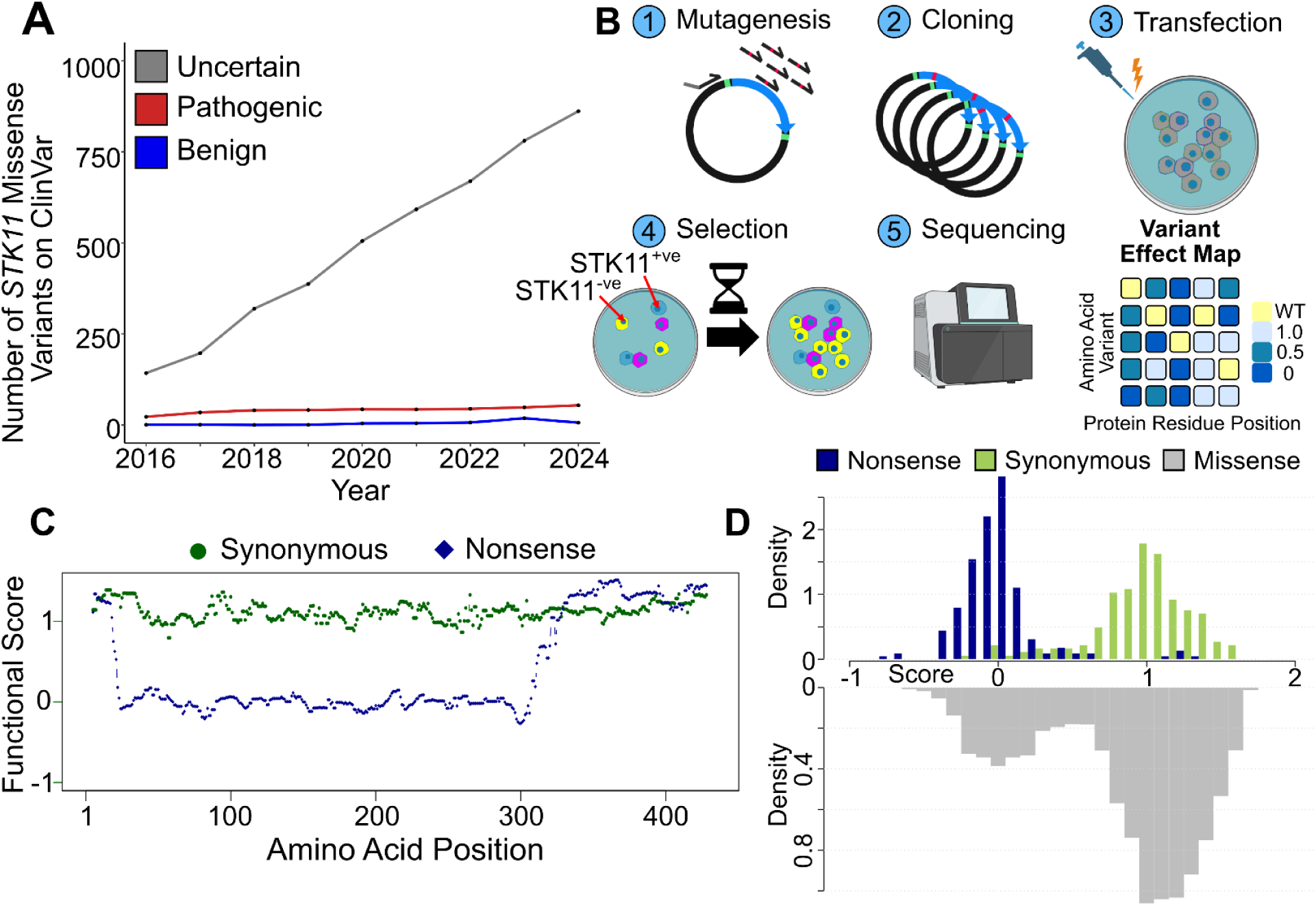
Distributions of *STK11* Functional Scores. **A** *STK11* missense variants and their associated clinical classifications were extracted from ClinVar for every month between January 2016 to December 2024, with the yearly average plotted on the x-axis. The number of unique missense variants in *STK11* is plotted on the y-axis with the blue line representing variants with likely benign or benign classifications, red indicating likely pathogenic or pathogenic, and gray for VUS or variants with conflicting interpretations. **B** Overview of the deep mutational scanning workflow applied to *STK11* (images of plasmids, cell culture, and sequencing generated via BioRender). **C** The median functional score for synonymous variants (green) and nonsense variants (blue) was calculated in a moving window analysis. Each window contained 10 amino acids and the analysis moved along *STK11* one position at a time. **D** Functional scores for *STK11* amino acids 22 to 311 were plotted as a histogram, separated into nonsense (blue), synonymous (green), and missense variants (gray).

### Quality control and initial evaluation of the STK11 variant effect map

After combining technical replicates, we filtered variants from each biological replicate to retain only those that were sufficiently represented pre-selection to be considered well measured, and removed variants exhibiting strong disagreement between replicates (see Methods). We next combined degenerate codons for remaining variants within each biological replicate, then examined agreement for each amino acid pairing of the three biological replicate scores. These showed correlation that was modest but highly significant (Pearson correlation coefficients ranging from 0.5 to 0.52; all p-values below 1×10^−308^) and were in agreement (see Methods) for over 80% of variants (Supplementary Fig. 2). After further filtering to remove variants for which biological replicates disagreed (see Methods), we combined biological replicates, yielding scores for 6,026 (∼73%) of the 8,208 possible amino acid substitutions (Supplementary Fig. 3). Of the 2,568 substitutions that are reachable by single-nucleotide change (and therefore more likely to be observed in humans), the combined map covered 2,049 (∼80%) variants. (Supplementary Table 1).

In addition to the 6,026 scored missense variants, the final combined *STK11* variant effect map contained similarly-quality-control-filtered functional scores for 287 synonymous and 339 nonsense variants. To evaluate the separation between scores of synonymous and nonsense variants, and to identify regions with reduced separation, we carried out a moving window analysis across the *STK11* protein sequence (Fig. 1c). Synonymous variants were nearly uniformly tolerated across the protein, as expected, with ∼94% of synonymous variants showing ‘tolerated’ (> 0.5) scores.

Between positions 22 and 311, our map showed ∼96% of nonsense variants to have damaging (below 0.5) scores (Fig. 1d) and discordant nonsense and synonymous variants (i.e., apparently-tolerated nonsense or apparently-damaging synonymous variants) did not appear to cluster at specific residue positions or regions. By contrast, at both the C- and N-termini, nonsense variants appeared largely tolerated. At the C-terminus, ∼95% of nonsense variants after position 311 had tolerated functional scores. This tolerant section of the C-terminus appears to begin after the end of the STK11 kinase domain (positions 49 to 309), suggesting that a large portion of the C-terminus is not required for cell growth. Of the first 22 amino acid positions at the N-terminus, Met22* and Gln7* are the only nonsense variants with a damaging (below 0.5) functional score. Met22 could serve as an alternative translation initiation site. The Gln7* result is difficult to reconcile with this model, but given that the 10 other well-measured stop codons from positions 1 to 21 (inclusive) appeared tolerated, we attribute the Gln7* result to experimental variation. Further support for the alternative start model comes not only from the fact that the Met22 context matches the known ‘GnnAUGG’ mammalian ribosome-binding motif^23^, but that the enrichment for truncating mutations that can be observed for somatic mutations in tumor samples is nearly absent in the first 21 amino acid positions of STK11 (Supplementary Fig. 4).

Missense variant scores followed a bimodal distribution with modes that roughly lined up with those of synonymous and nonsense variants (Fig. 1d). Out of 6,026 missense variants with functional scores across the entire protein, 993 (16.5%) had scores below 0.5 (non-functional) and 5,033 scored above 0.5 (functional). Thus, the majority of *STK11* missense variants exhibited suppressed growth in the HeLa proliferation assay (i.e., were tolerated) while a subset of missense variants retained a growth advantage (i.e., were damaging). The fraction of variants that appeared damaging rose (to 23.5%) when subset to positions 22-311.

### Mutational hotspots at known kinase motifs are intolerant to variation

Protein kinases often share conserved amino acid sequence motifs corresponding to functions necessary for kinase activation and phosphorylation of the substrate. Several STK11 kinase motifs have also been identified as mutational hotspots in pan-cancer tumour sequencing surveys^24,25^. Describing these STK11 motifs in N- to C-terminal order: GxGxxG motif residues (matching STK11 positions 56-61) form the flexible glycine rich loop needed for orienting ATP; VAIK motif residues (matching RAVK at STK11 positions 75-78) form a salt bridge with a conserved glutamic acid (at position 98 in STK11); HRD motif residues (matching HKD at STK11 positions 174 to 176) and a conserved asparagine (at position 181 in STK11) form the active site of the catalytic loop; DFG motif residues (DLG at STK11 positions 194 to 196) serve to trigger a conformational change during activation; and finally APE motif residues (STK11 PPE at positions 221-223) stabilize the C-lobe of the kinase.

For conserved positions at each motif, median scores were significantly lower than the median over all non-motif positions in the kinase domain (median 1.04), by Wilcoxon rank-sum tests: GxGxxG (median = 0.23; p =2.4×10^−8^), VAIK (median = 0.22; p =4.4×10^−5^), salt-bridging-glutamic acid (median = 0.29; p =1.6×10^−3^), HRD motif (median = 0.23; p =2.4×10^−8^), position N181 (median = −0.04; p =2.89×10^−10^), and DFG (median = 0.14; p =2.4×10^−3^), and APE (median = 0.67; p =6.7×10^−5^) (Fig. 2A). Where STK11 elements corresponding to motifs differed from the consensus motif sequence (VAIK, DFG, HRD, and APE), restricting to positions matching the consensus sequence further decreased the motif median (VAIK Δscore = -.23; DFG Δscore = −0.33; HRD Δscore = −0.17) except for APE (Δscore = 0.06) (Supplementary Fig. 5).

**Figure 2.**
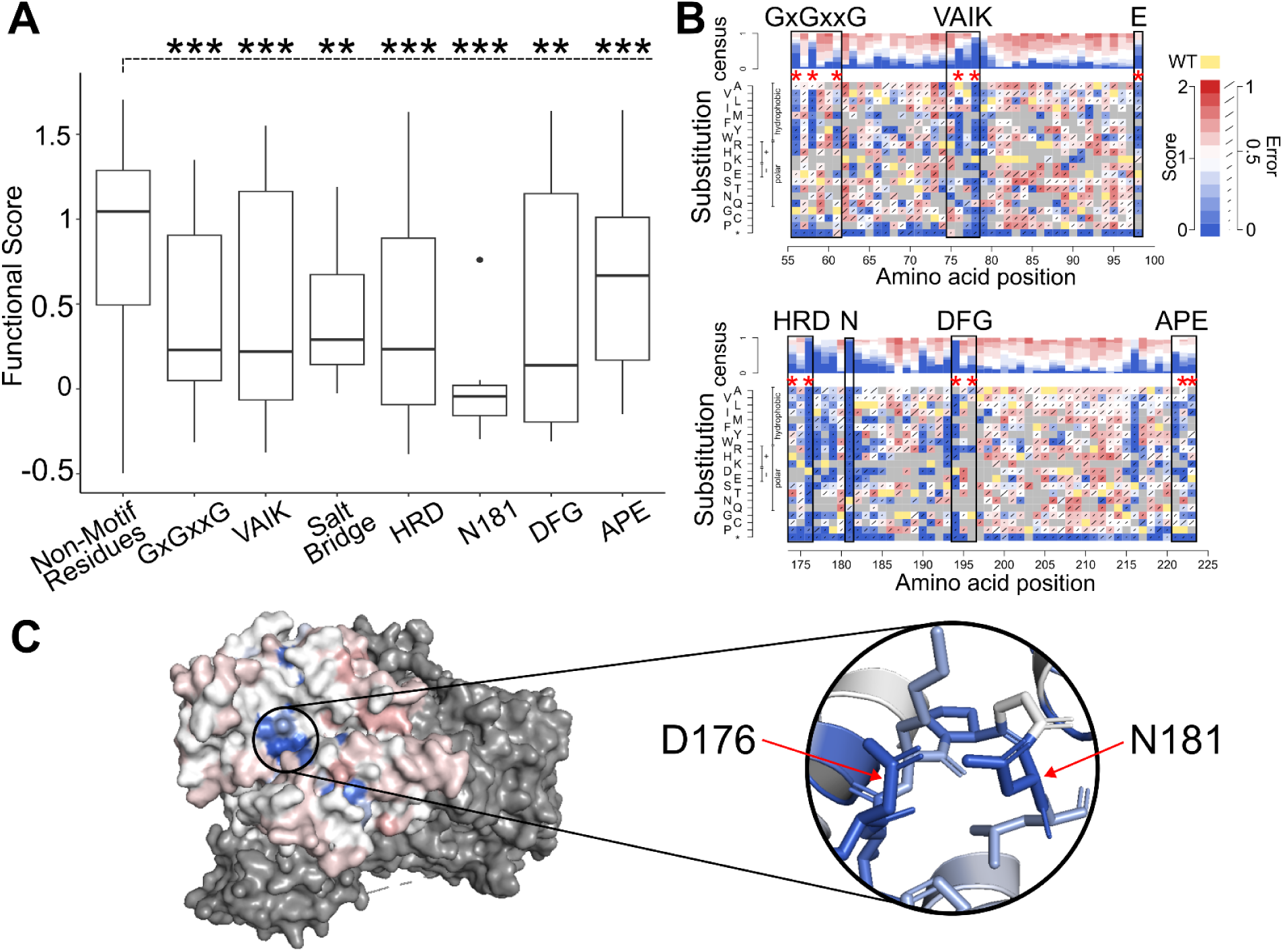
Protein Motifs Corresponding to Known STK11 Functions are Intolerant to Variation. **A** Mutational hotspots identified in Hudson *et al.* were separated into distinct kinase activation motifs (GxGxxG, VAIK, Salt bridge, HRD, DFG, and APE) and plotted against *STK11* functional scores for each variant at these positions. Boxplots depict the median functional score for each motif as well as the 25th and 75th percentiles with vertical lines extending to 1.5 times the interquartile range above and below 25th and 75th percentiles. Significant differences from non-hotspot positions (Mann Whitney U test) are indicated by double star (P<0.01) or triple star (P<0.001). **B** *STK11* variant effect map positions corresponding to the kinase motifs are shown, where x-axis is the position along *STK11*, y-axis is the substituted variant, and the color indicates functional score (damaging scores shown in blue, tolerated in white/red, variants that did not pass filtering in gray, and the wild-type residue in yellow). Error rates, calculated as described in the methods, are indicated by increasingly long diagonal lines for each variant. **C** The median map score at each position was ‘painted’ onto the *STK11* structure (PDB: 2WTK, residues 43 to 347), where dark blue indicates damaging scores less than 0.2, successively lighter shades of blue for scores between 0.2 and 0.8, white for tolerated scores between 0.8 and 1.2, and red for scores above 1.2. Dark gray represents STK11’s binding partners, MO25 and STRAD. Both the surface level view and a focused look at the active site are provided.

Closer inspection of the variant effect map showed additional biological insights. Substitutions at specific motif positions — the last two conserved G positions (G58 and G61) of GxGxxG, K76 in VAIK, and the salt bridging E98 — all showed hydrophobic substitutions to be more damaging than polar substitutions (Δmedians −0.99, −0.56, −0.16, and −0.32 at positions 58, 61, 76 and 98, respectively). (Fig 2B). Almost all substitutions were damaging at K78 in the VAIK motif, position D176 in the HRD motif, at the active site asparagine N181, and at D194 in the DFG motif. Previous reports suggested that K78 is directly involved in salt bridge formation, D176 accepts a hydrogen removed from the substrate during phosphorylation, and D194 chelates magnesium ions^24,26^. Interestingly, despite measuring all 19 possible missense variants at position N181, substitution to glutamine was the only variant receiving a tolerated score of 0.76, suggesting the importance of both the size and charge characteristics that are shared between asparagine and glutamine residues (Fig 2C).

### Integrating estimates of protein stability, conservation, and solvent accessibility

We compared functional scores at each position with other characteristics: predicted protein stability (ΔΔG), conservation, and solvent accessibility of the wild-type residue.^27^ Functional scores were summarized in terms of median score at each position, and other characteristics were derived as follows: a position was annotated as 1) “instability-prone” if the median predicted ΔΔG of substitutions was greater than 1; 2) “conserved” if the ConSurf score was below the published default threshold of −0.467; 3) and “buried” if the side-chain accessibility was less than 20% of the maximum possible for that side chain^28–30^. A comparison between functional score and these characteristics of the STK11 protein sequence (Fig. 3A) confirmed expected features and provided clues about mechanisms underlying variant dysfunction.

**Figure 3.**
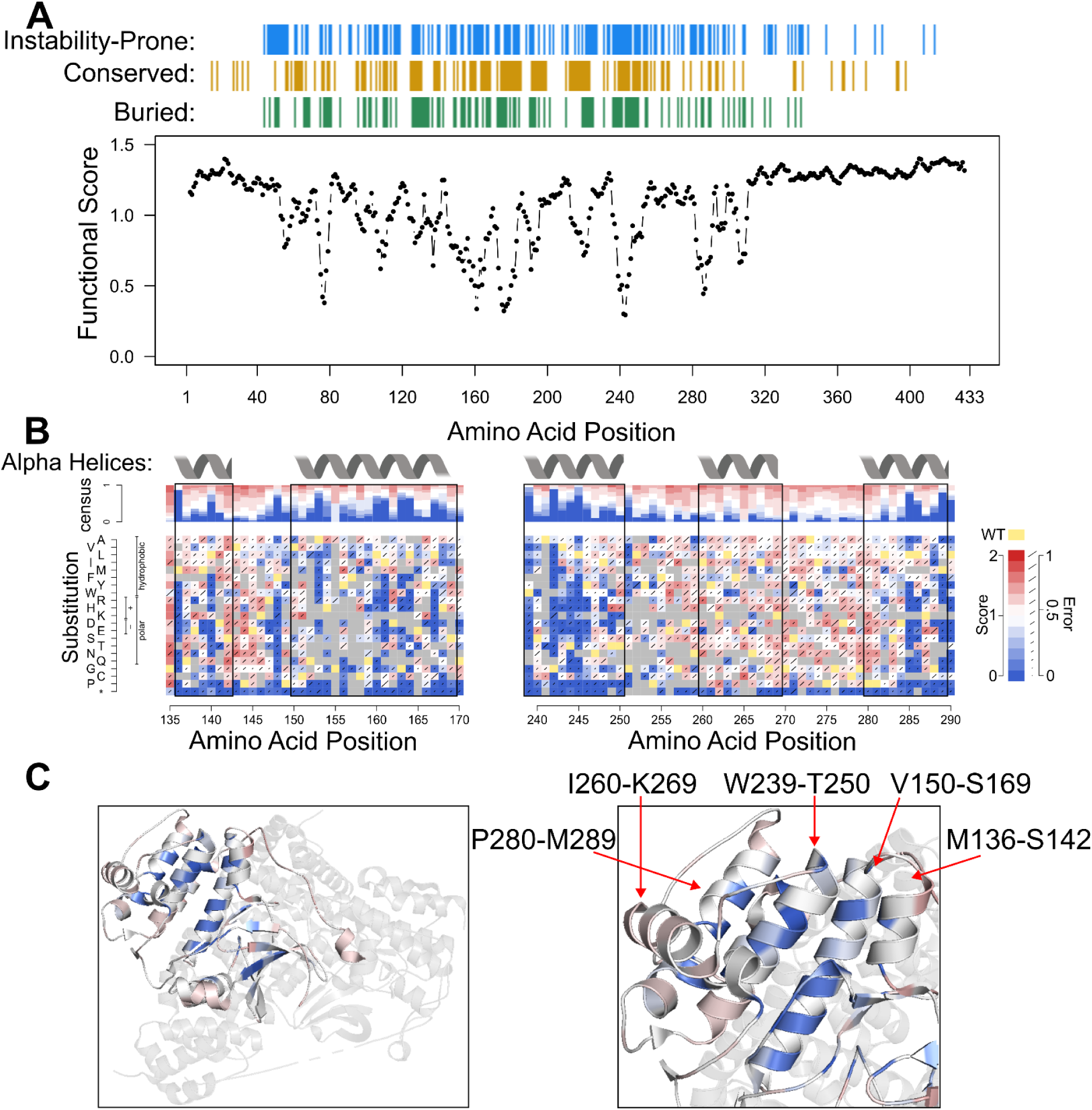
Moving Window Analysis of Variant Impacts on Function, Protein Stability, Conservation, & Solvent Accessibility. **A** The median functional score for substitutions in each 10-amino acid wide moving window (shifting to each position in turn). Positions with predicted ΔΔG suggesting instability (median ΔΔG greater than 1) are indicated on the horizontal blue track above the plot, where ΔΔG was calculated with FoldX (v.4.0) using an STK11 PDB structure predicted by AlphaFold v.2022-11-01, Monomer v.2.0 pipeline^31^). Positions found to be conserved according to multiple sequence alignments on the ConSurf web server (ConSurf scores below −0.467) are shown in gold, and positions with low relative solvent accessible surface area below 20% side-chain accessibility (FreeSASA software v.2.02, October 22, 2017) are labeled on the green horizontal track. **B** Partial variant effect maps are included for segments described in the main text (positions 136 to 169 and 239 to 289) with labels and black boxes for alpha helices found on *STK11* UniProt entry Q15831. **C** Median map scores were ‘painted’ onto STK11 (PDB: 2WTK, residues 43 to 347) as described in Figure 2C. Secondary structures (alpha helices, beta sheets) are depicted with notations for alpha helices at positions 136-142, 150-169, 239-250, 260-269, and 280-289.

First, this analysis revealed that the terminal regions tended to tolerate missense variants, consistent with their tolerance of nonsense variants as described above. The comparison with other sequence features for the tolerated terminal regions (defined by eye for missense variants as the N-terminus before position 40 and the C-terminus after position 340) also showed few (6.9%) instability-prone or buried (0.8%) positions, but some (19.0%) conserved positions. A more quantitative analysis found both scores and characteristics to be significantly different in these terminal regions from non-terminal positions, confirming that these positions were more tolerated (Δmedian functional score = 0.21; p =3.3×10^−29^), more stable (Δmedian ΔΔG = −1.21; p =1.78×10^−21^), less conserved (Δmedian ConSurf score= 0.539; p =1.87×10^−5^), and more solvent-exposed (Δmedian solvent accessibility = 63.6%; p =4.12×10^−40^).

Second, the analysis was consistent with the functionality of the kinase motifs discussed above. More specifically, it showed that 56.2% of motif positions were instability-prone, 94.1% were buried, and 84.6% were conserved (in contrast to 38.6%, 34.9%, and 28.0%, respectively, for non-motif positions). Relative to non-motif positions, motif positions were also quantitatively more likely to be instability-prone, conserved, and buried than non-motif positions with (p = 0.01, 1.8×10^−8^, and 2.7×10^−5^, respectively; one-tailed Wilcoxon).

The sensitivity to variation we observed within alpha helices tended to be consistent with their location in the 3-D crystal structure (PDB: 2WTK). For example, two annotated alpha helices (positions 136-142 and 150-169) contained distinct positions (136, 140, 148, 153, 160, 161, 163, 164, and 167) with damaging median functional scores (Fig. 3B). Comparison with other characteristics showed 100% of these positions to be instability-prone, 77.8% conserved, and 100% buried. All other positions between 136 and 169 showed tolerated median functional scores (overall median = 1.06), with fewer (47.8%) predicted to be instability-prone, fewer (48%) conserved, and fewer (37.5%) buried. ‘Painting’ functional scores onto the 2WTK *STK11* crystal structure revealed patterns of tolerance in which the side of the helix that is buried (either within the STK11 structure or by MO25-STRAD binding partners) or facing the active site were more sensitive to variation, while surface-exposed faces tolerated substitution (Fig. 3C). Similar findings hold true for alpha helices at positions 239-250, and 280-289, where positions with damaging functional scores were instability-prone, conserved, buried, and coincided with internal-facing regions. Interestingly, the alpha helix at position 260-269 is entirely external to the core of *STK11* and no position was observed to be intolerant to substitution (Fig. 3C). Taken together, alpha helices 136-142, 150-169, 239-250, and 280-289, show patterns of tolerance that are consistent with their positions and collective role in surrounding and shaping the active site.

### Comparisons with alternative functional assays

Previous studies have characterized functional impacts of *STK11* variants by measuring either STK11 substrate phosphorylation or cellular growth rate in various model systems^24,32–34^. Our map scores were strongly correlated with a study of the effects of 20 STK11 variants on p53 transcriptional activation in A549 cells, (PCC = 0.92; *P* = 4.5×10^−8^; one-tailed Wilcoxon) (Fig. 4A). Variants with damaging map scores displayed lower levels of p53 transcriptional activation (median 0.51) than those with tolerated map scores (median 1.17; |Δ median| = 0.66; *P* = 3.1×10^−5^; one-tailed Wilcoxon).

**Figure 4.**
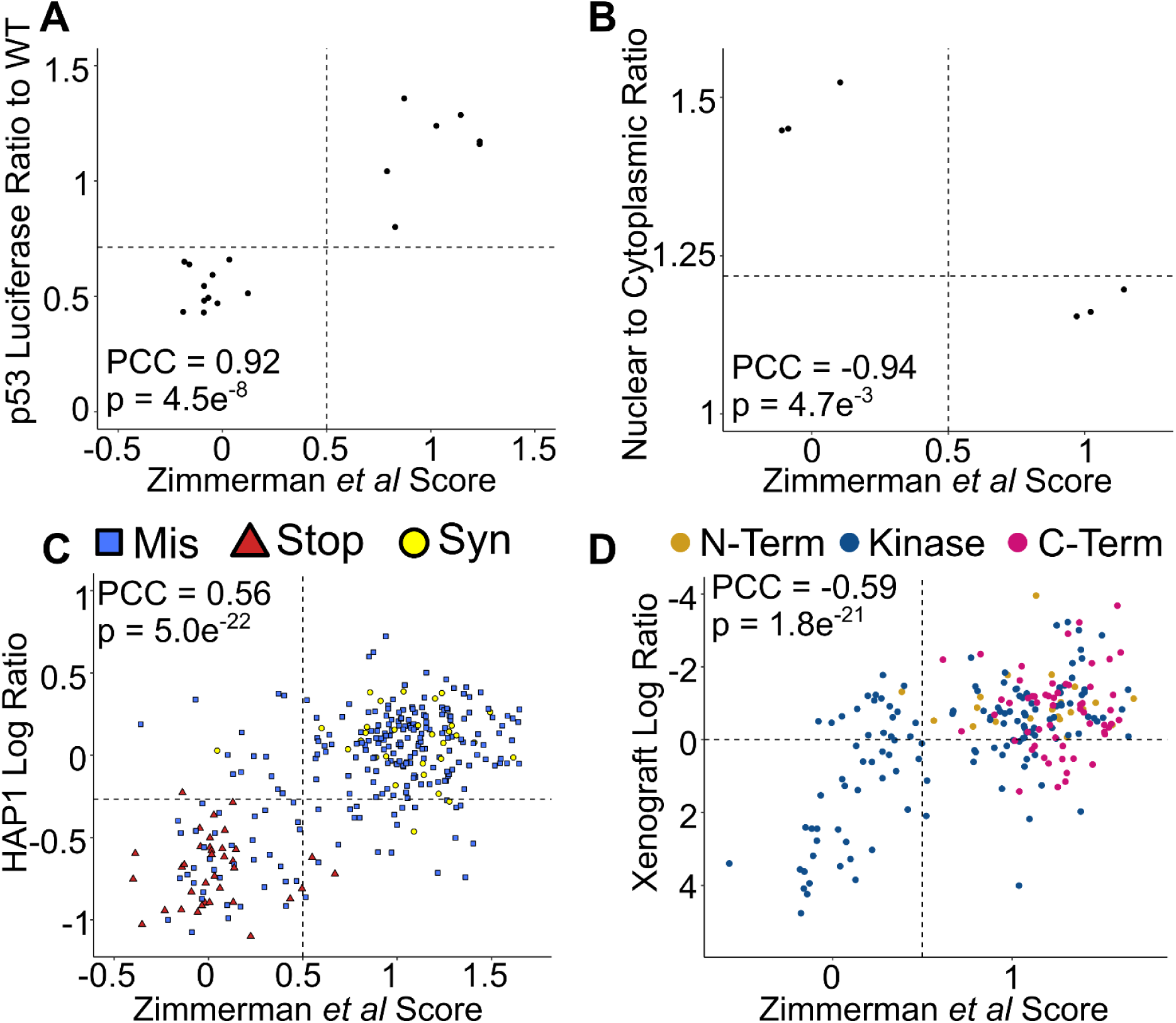
Variant Effect Map Scores Correlate with Alternative Functional Assays. **A** Our map scores were compared with results from a p53 luciferase based reporter system^38^. For each variant, P53 luciferase activity was plotted relative to expression induced by wild-type *STK11*. Intermediate scores are indicated by horizontal and vertical dashed lines. **B** Our map scores were compared with measurements of STK11 nuclear localization in HeLa cells. Here, *STK11* variants were fused to GFP and expressed in HeLa cells via a lentiviral vector. Cells were analyzed by fluorescence microscopy and the relative GFP intensity was calculated between the cytoplasm and nucleus. The nuclear to cytoplasmic ratio of GFP for each variant was compared with our map score. An intermediate proliferation score is indicated by the dashed vertical line and the wild-type STK11 nuclear/cytoplasmic GFP ratio indicated by the horizontal dashed line. **C** Our map scores were compared with functional scores measured via saturated genome editing^36^ of *STK11* exon 5 in HAP1 cells. Loss of *STK11* function in HAP1 cells reduces their ability to proliferate and survive, leading to drop-out in the pooled population. Missense variants are shown as blue squares, nonsense variants as red triangles, and synonymous variants as yellow circles. **D** Our map scores were compared with the impact of STK11 variants on tumor growth of A549 cells xenografted into mouse (expression of wild-type STK11 reduces growth of tumors formed by these A549 cells, so that A549 cells carrying reduced-function STK11 variants are enriched in mouse tumors).

To enable further comparisons, we generated three additional datasets using diverse model systems and phenotypic readouts. First, motivated by the knowledge that activated STK11 shuttles to the cytoplasm, we examined the nuclear-cytoplasmic localization of STK11 variants in HeLa cells using an established protocol^35^. For a set of 6 variants (3 arbitrarily-chosen pathogenic and 3 benign), we found our map scores to be strongly anti-correlated with the nuclear-cytoplasmic ratio of STK11-GFP (PCC = −0.94; *P* = 4.7×10^−3^; one-tailed Wilcoxon) (Fig. 4B), as expected. Variants with damaging scores in our map showed higher levels of STK11 nuclear accumulation (median 1.45) than those with tolerated map scores (median 1.16; |Δ median| = 0.29; *P* = 0.05; one-tailed Wilcoxon).

Second, we performed assays in an independent cell model, using HAP1 cells in which loss of *STK11* function reduces cellular survival. We applied saturation genome editing (SGE)^36^, generating mutagenesis libraries for exons 5 and 9 (which spans positions 200 to 245) and exon 9 (which spans positions 370 to 433) and, in separate experiments, integrating these libraries at their respective endogenous loci via CRISPR-Cas9 stimulated homologous recombination. Relative abundance of variants was measured before and after 21 days of growth. In total, scores were obtained for 702 missense variants (290 in exon 5, 412 in exon 9), of which 587 also had map scores (244 in exon 5, 343 in exon 9). Map scores were significantly and positively correlated with HAP1 growth in exon 5 (PCC = 0.56, *P* = 4.95×10^−22^; one-tailed Wilcoxon). By contrast, we observed no correlation between our map and SGE scores in exon 9 (PCC = −0.02, *P* = 0.67; one-tailed Wilcoxon), which can be explained by the dearth of variation in this region in both our map and the SGE data (most variants were tolerated), limiting our ability to observe co-variation (Fig. 4C) (Supplementary Fig. 6). The HAP1 essentiality assay may, however, have had greater sensitivity to damaging variation in this region than our HeLa toxicity assay; for example, of the 7 variants measured at positions both Q382 and R383, three at each position were found damaging in the HAP1 assay (in contrast to the HeLa assay, which found 0 damaging variants despite well-measured scores for 18 and 19 variants at positions Q382 and R383 respectively).

Third, we tested the function of *STK11* variants by introducing variant-expressing A549 as mouse xenografts, in which expression of wild-type STK11 restricted growth of xenografted tumor cells. More specifically, A549 cells expressing a pool of barcoded *STK11* variant cDNAs were implanted into mice, with variants quantified at the end of the study via barcode sequencing. The 215 missense variants with functional scores in both the xenograft assays and our HeLa proliferation-based map showed a significant negative correlation, as expected (PCC = −0.59, *P* = 1.76×10^−21^; one-tailed Wilcoxon) (Fig. 4D). While nearly all variants that our map scores had found to be damaging in the HeLa assay were located in the kinase domain (between positions 49 to 309), variants found to be damaging by the xenograft assay were split, with 52 variants found in the kinase domain and 12 variants in the C-terminus (past position 309). Thus, the xenograft assay may be more able to identify C-terminal STK11 variant functional impacts than our HeLa toxicity assay. Examples include: T363N (located at a known DNA-damage associated phosphorylation site); R383C and R384W (positions also found to be important in the HAP1 assay); and R409W, located in a membrane localization motif (positions 403 to 426) and known to cause nuclear sequestration of STK11 while retaining kinase activity^37,38^. Each of these variants showed enrichment in tumors over time (log_2_-ratios of 0.27, 0.50, 0.91 and 1.15 respectively).

### STK11 variant effect assays distinguish pathogenic from benign variation

A key question about our HeLa toxicity-based *STK11* variant effect map is whether it can distinguish clinically annotated pathogenic variants from benign. To evaluate the ability of these scores to separate pathogenic from benign variants, a reference set consisting of 20 pathogenic or likely pathogenic and 109 benign or likely benign missense variants was assembled by Ambry Genetics (author APLM). We first used this reference set (or at least the subset of assayed reference variants) to evaluate the subset of the above-described smaller-scale functional assays for which at least 2 pathogenic and benign variants had been assayed. This evaluation was in terms of both precision (fraction of variants with scores below a given threshold that are known to be pathogenic) and recall (fraction of known pathogenic variants that received a score below this threshold). Because precision is dependent on the (somewhat arbitrary) balance of pathogenic and benign variants in the reference set, we instead calculated “balanced precision” --the precision expected if the reference sets had contained an equal number of pathogenic and benign variants. Each of the small-scale assays performed well: 1) the HAP1 essentiality assay achieved a near perfect Area Under the Balanced Precision-Recall Curve (AUBPRC) of 0.98 and Recall at 90% Balanced Precision (R90BP) of 1.00 (based on 2 pathogenic and 53 benign reference variants); 2) the A549 Tango xenograft assay reached an AUBPRC of 0.92 and R90BP of 0.71 (based on 7 pathogenic and 20 benign reference variants). (See Supplementary Fig. 7).

Before applying the same evaluation to our larger-scale HeLa-toxicity scores, we sought to provide a subset of only the highest-quality functional scores that would be more appropriate for use in clinical variant interpretation. To this end, we removed variants with a confidence interval overlapping the intermediate score of 0.5 (see methods), leaving 5,334 missense variants with high-confidence scores. Of these, 11 pathogenic and 75 benign reference variants had a high-confidence score. The *STK11* variant effect map performed well in distinguishing pathogenic from benign variants, achieving an AUBPRC of 1.00 and capturing 100% of pathogenic variants at a stringency of 90% Balanced Precision (i.e., 100% R90BP) (Fig. 5A). Because performance without the confidence interval filter was slightly reduced (AUBPRC 0.99 and R90BP 0.92) (Supplementary Fig. 7), we used only post-filter scores from our map for subsequent clinical evidence calibration.

**Figure 5.**
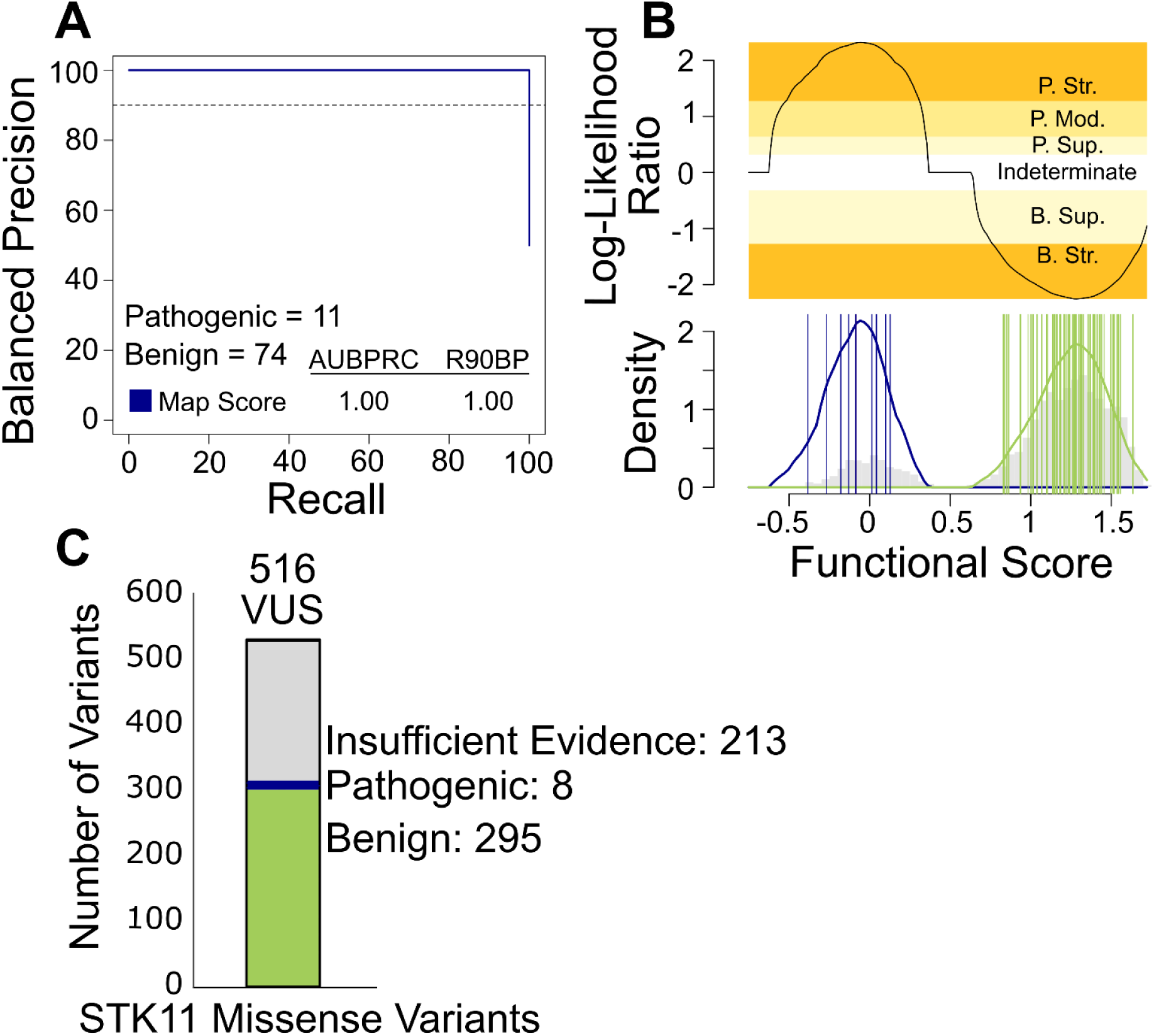
*STK11* Functional Scores Accurately Classify Clinical Variants. **A** Using our *STK11* map scores and a known set of clinically annotated pathogenic or benign *STK11* variants from Ambry Genetics, we evaluated balanced precision—defined at each score threshold by the fraction of variants that are pathogenic given a test set that is balanced (i.e. has a 50% prior probability of pathogenicity)—versus recall (fraction of pathogenic variants captured at this threshold). The horizontal dashed line indicates recall and 90% balanced precision (R90BP) with the numerical area under the balanced precision vs. recall curve (AUBPRC) and R90BP listed in the bottom-left hand legend. **B** LLRs of pathogenicity were calculated by comparing the probability distributions of scores for known pathogenic and benign variants in the reference set. The log ratio of the likelihood of observing a score in the positive pathogenic reference set (blue) compared to the negative benign reference set (green) was calibrated to ACMG evidence strengths. Probability distributions are overlaid on the gray histogram of *STK11* missense variant scores with the top panel showing which score ranges correspond to each ACMG evidence strength. Evidence towards pathogenicity or benignity are split into discrete categories of strong (Str.), moderate (Mod.), and supporting (Sup.). **C** The pathogenicity of a set of *STK11* VUS variants that had been previously classified by Ambry Genetics were re-evaluated, using ACMG evidence strength values derived from our calibrated functional data together with all other available evidence. The stacked bar chart indicates the number of VUS with sufficient evidence for reclassification towards pathogenicity (blue), benignity (red), as well as those remaining as VUS (gray).

For the purposes of assisting with variant interpretation, *STK11* functional scores were calibrated to differing levels of ACMG evidence strength. To this end, the Ambry reference set was used to estimate, separately for pathogenic and benign reference variants, a continuously-valued score distribution. Using these distributions, the relative likelihood of pathogenicity can be calculated for any given score of interest. Using this approach, we calculated a Log-Likelihood Ratio of pathogenicity (LLR) value for every variant. These LLR values were converted to the categorical system of evidence strengths described in ACMG/AMP guidelines using an adaptation^39^ of the original Tavtigian et al. approach^40^. Thus, an LLR value above 2.5 indicates “very strong” evidence of pathogenicity; LLR between 2.5 and 1.3 indicates “strong” evidence; LLR from 1.3 to 0.64 indicates “moderate” evidence, and LLR from 0.64 to 0.32 provides “supporting” evidence of pathogenicity. Negative LLR values between −0.32 and −1.32 and LLR values below −1.32 indicate “supporting” and “strong” evidence of benignity, respectively. Using this calibration, 1, 4, and 17 of the *STK11* variants that had previously been classified as VUS by Ambry, received new supporting, moderate, and strong evidence of pathogenicity, respectively from our STK11 map. Moreover, an additional 6 and 322 Ambry VUS received supporting and strong evidence of benignity, respectively (Fig. 5B). Thus, the map provided new functional evidence for 350 (68%) of the 516 *STK11* missense variants that Ambry had previously classified as VUS.

The Ambry Genetics team (authors APLM, FH, MER) followed their formal variant curation process to predict the results of re-evaluating all 516 previous VUS variants, integrating *STK11* map scores with additional ACMG-recognized sources of evidence. As a result, 8 would be reclassified towards pathogenic (4 likely pathogenic and 4 pathogenic) and 295 would be reclassified towards benign (139 likely benign and 156 benign) (Fig. 5C).

### Quantitatively relating functional scores to PJS disease phenotypes

Next, we sought to go beyond variant classification and quantitatively assess the correspondence of our map scores with patient phenotypes. To this end, we extracted de-identified phenotypic information, including age at first diagnosis of PJS or of a PJS-related cancer, from both PJS-related publications and for a PJS cohort (Mt Sinai Hospital, Toronto, Canada; Research Ethics Board REB # 23-0006-C)^41–51^. Map scores showed a significant positive correlation with age at PJS diagnosis (PCC = 0.52, *P* = 3.6×10^−3^) and age at PJS-cancer diagnosis (PCC = 0.94, *P* = 1.7×10^−2^) (Fig. 6A).

**Figure 6.**
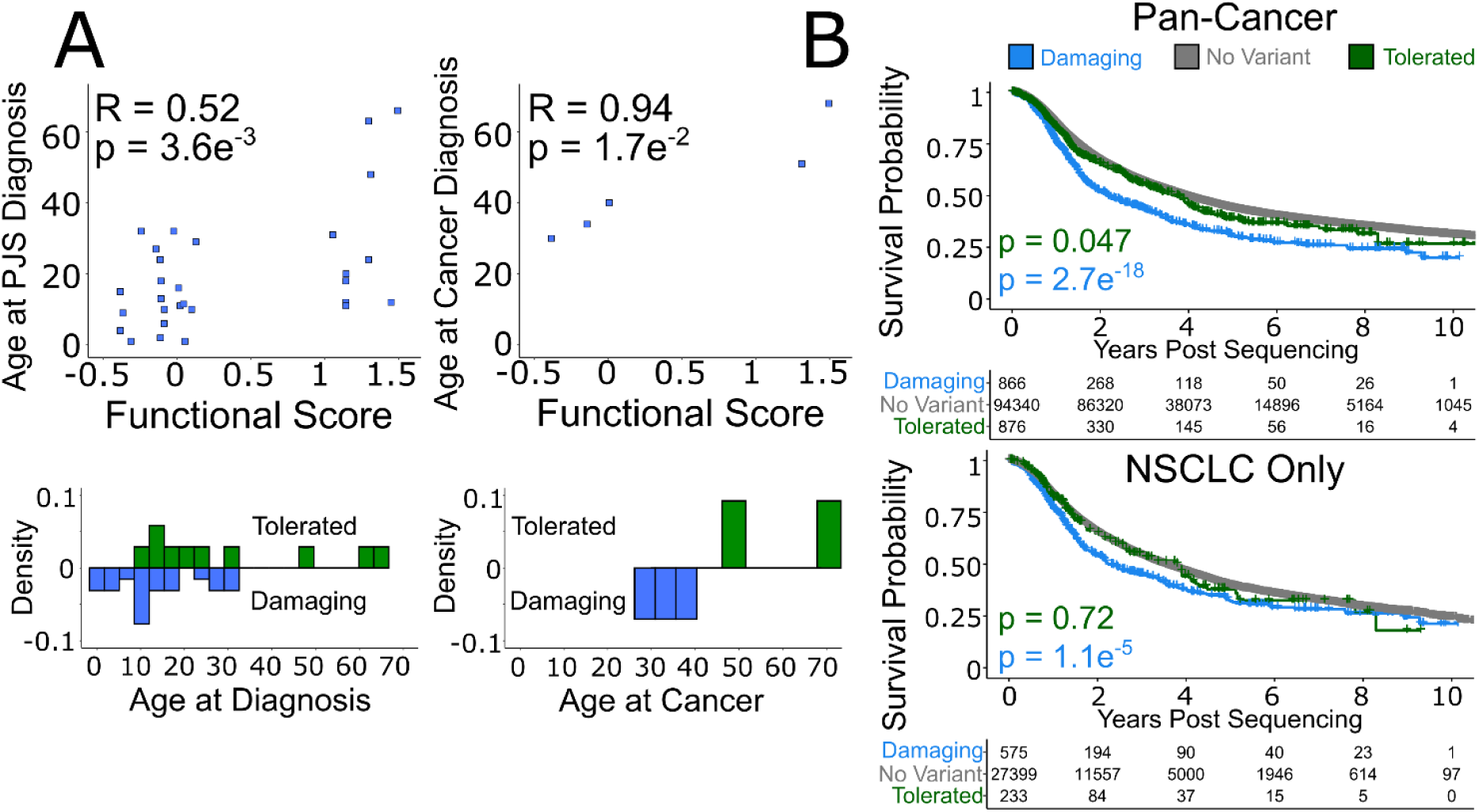
Detrimental Functional Scores Are Associated with Earlier Diagnosis of PJS and Onset of Cancer. **A** Age at initial PJS diagnosis and age at incidence of cancer for individuals previously diagnosed with PJS were collected from patient charts at the Zane Cohen Centre as well as from a literature review of available PJS case studies. The age at first diagnosis often coincides with the earliest manifestation of symptoms (such as hyper-pigmentation, intussusception, etc.). Missense variants are plotted as blue squares. Pearson correlation between age at PJS diagnosis or age at PJS-associated cancer and functional score is provided. **B** Metrics relating to overall survival for individuals with cancer were extracted from project GENIE (as of March 2025). We collected age at which sequencing was reported, age at last contact, and age at death if applicable. Patients were stratified into those without an *STK11* variant (gray), a tolerated *STK11* variant (green), or a damaging *STK11* variant (blue) according to our map score. Kaplan-Meier survival curves visualized the probability of survival over time in years, each ‘drop’ representing a death end-point, and each ‘tick’ a currently living individual. Log-rank tests were performed to compare outcomes of individuals carrying damaging, tolerated or no *STK11* variant.

Patients carrying missense variants with damaging map scores were associated with an earlier age of diagnosis than those with tolerated variants for both PJS (median age 11.5 versus 22; *P* = 1.2×10^−2^, one-tailed Wilcoxon) and non-significantly for PJS-related cancer (median age 34 vs 59.5 years; *P* = 0.1, one-tailed Wilcoxon). Damaging missense variants were statistically indistinguishable from nonsense variants in terms of age at PJS diagnosis and age at PJS-cancer diagnosis (|Δ median| = 4.5, 2 years; *P* = 0.50, 0.2; one-tailed Wilcoxon). (See Supplementary Fig 8).

### Quantitatively relating functional scores to non-PJS-related cancer phenotypes

To investigate potential quantitative relationships between our functional scores and cancers with somatic, as opposed to germline, mutations in STK11, we collected information relating to cancer patient survival from a non-redundant set of clinical studies on cBioPortal, as well as from project GENIE^52,53^. In project GENIE, 2,836 (∼1.3%) individuals had tumours bearing a *STK11* missense variant, 1,926 of which were scored in the map. Aggregating information across all cancer types found that patients with an *STK11* missense mutation scoring as tolerated in our map showed survival that was nearly statistically indistinguishable from individuals without a *STK11* variant (times to 50% survival of 3.72 vs 3.89 years; P = 0.047, log-rank test) (Fig. 6B). Alternatively, patients whose tumors carried STK11 mutations that were scored as damaging in our map exhibited higher mortality than those without a variant (time to 50% survival 2.17 years vs 3.89 years; P = 2.7×10^−18^, log-rank test). Similar findings hold true if we subset to NSCLC, where tolerated variants were statistically indistinguishable from individuals without a *STK11* variant (times to 50% survival of 3.80 vs 3.48 years; P = 0.72, log-rank test), while damaging variants exhibited higher mortality than those without a variant (time to 50% survival 2.26 years vs 3.48 years; P = 2.7×10^−18^, log-rank test) (Fig. 6B). These findings were confirmed using the curated non-redundant set of clinical studies hosted on cBioPortal (Supplementary Fig 9).

## Discussion

Here we applied a mammalian cell-based assay for *STK11* variant effects at scale. The resulting map was largely concordant with smaller-scale functional assays, and also recapitulated many known biochemical features of STK11. The map also provided some interesting new molecular-genetic or biochemical observations, including an alternative translation initiation site at residue Met22, and a refined understanding of which positions within and surrounding the active site are most sensitive to variation. As discussed below, our map holds potential clinical value both for diagnosing and informing therapy for those with PJS, and also for those with NSCLC or other tumours bearing *STK11* variants.

Because the intestinal polyps caused by PJS contribute to incidences of bleeding, intestinal blockage, and intussusception (i.e. improper intestinal folding), and the increased lifetime risk of cancer, it is currently recommended that those diagnosed with PJS: 1) undergo screening for intestinal polyps (i.e. colonoscopy, endoscopy) starting at eight years old, with pre-emptive surgical removal of large polyps to prevent intussusception; 2) be screened for breast and gynecological cancers starting at the age of 25; and 3) be considered eligible for pancreatic cancer screening regardless of family history^54^. As a germline condition for which nearly all cases have been attributed to the *STK11* locus, genetic testing offers a potentially gold-standard method for diagnosis, but this potential is currently limited by the fact that 94% of clinical missense variants in *STK11* are currently classified as VUS or have conflicting classifications.

After applying filters to retain only the highest quality scores that would be most suitable for potential clinical use, our map covered ∼65% (5,334) of all possible amino acid changes and ∼68% (1,737) of amino acid changes reachable by single-nucleotide changes. Moreover, it provided at least supporting evidence for variant classification for ∼65% (5,318) of all missense variants.

Of the missense variants for which our map provided classification evidence, 350 had been previously encountered and classified as VUS by Ambry Genetics. When subjected to re-evaluation in the context of ACMG/AMP guidelines, 303 (59%) of these 516 changed from VUS to a more definitive classification. Although many *STK11* missense variants have yet to be observed clinically, it is likely that every single possible nucleotide change within *STK11* already exists in someone alive today^11^. Thus, our proactive measurements will provide more rapid and definitive assessments for a large fraction of missense *STK11* variants, as they are observed in the future. Thus, proactive variant interpretation has the potential to limit unnecessary screening of individuals with benign *STK11* variants, while more confidently indicating monitoring for those carrying pathogenic PJS-causing variants.

In the context of the diverse cancer types in which *STK11* somatic variants are found, our map offers the potential to distinguish those in which *STK11* is a passenger mutation as opposed to being a cause (driver) of the cancer. This is potentially clinically actionable, in that driver *STK11* mutations are known to confer resistance to now-standard immunotherapies^2,3,55^. There is also some potential for cancers driven by *STK11* mutations to be more sensitive to alternative therapeutics^56–59^. For example, the potential to restore immunotherapeutic sensitivity to *STK11*-driven cancers by inhibiting CoREST deacetylase activity has been explored via Tango Therapeutics’ compound TNG260^60^, or via inhibitors of AXL, CDK4/6, or STAT3^61–63^. The fact that *STK11* mutation scores stratified patients in terms of survival outcomes suggests that our map could prove valuable in distinguishing oncogenic *STK11* driver mutations from passengers, with potential to improve outcomes through appropriately tailored therapies.

Our study had limitations, the most notable being that our assay appeared not to capture the impact of variants affecting regulatory post-translational modifications (PTMs) at the C-terminus. A likely explanation for this is that the STK11-toxicity activity we relied upon in our growth assay is not substantially altered by the presence or absence of these PTMs, despite their importance for STK11 localization and cellular polarity phenotypes. Other limitations stem from our evaluation of STK11 variants within cDNA constructs, as opposed to edits at the endogenous locus which might also capture effects that depend on the presence of introns (this most obviously includes splicing efficiency but also includes impacts on nonsense-mediated decay). A related limitation is that we did not ensure that STK11 was expressed at physiological levels, so that hypomorphic missense variants that only modestly lower STK11 activity might still reduce cell growth at our (likely higher than physiological) STK11 expression level and therefore be categorized as tolerated. Finally, there are many potential sources of random experimental error (e.g. base-calling error). Concerns about the impact of these limitations on the use of our map for separating pathogenic from benign variants are somewhat ameliorated by the fact that (after the stringent quality-filtering steps that we imposed) our map achieved perfect separation of previously-classified pathogenic and benign variants.

In conclusion, the *STK11* mutational scan presented here both strengthens our understanding of sequence-structure-function relationships and assists with variant interpretation. Resolving VUS to pathogenic or benign will both enable a non-invasive means of diagnosing PJS via a blood test and also support a genotype-driven approach to therapy for cancers involving somatic *STK11* variation.

## Materials and methods

### Plasmids & genomic integration

For the small-scale assay, a Gateway Entry (pDONR223) plasmid carrying the *STK11* open reading frame (ORF), corresponding to UniprotKB accession Q15831, was obtained from the Human ORFeome v8.1 library^64^. For the large-scale high-throughput assay, the *STK11* ORF was ordered from TWIST bioscience in the pTwist ENTR vector with an altered codon optimized DNA sequence to enable mutagenesis in GC-rich regions. The *STK11* ORFs were cloned into destination vectors for transfection into a HeLa cell line engineered to carry a ‘landing pad’ within the AAVS1 safe-harbour locus^19^, at which an EF1ɑ promoter is followed by a Bxb1 recombination site and an ORF encoding blue fluorescent protein (BFP). Destination plasmids contained a single Bxb1 recombination site upstream of the ORF, such that co-transfection with Bxb1 integrase can result in stable integration of the plasmid at the landing pad’s Bxb1 site, which not also disrupts BFP expression but also splits the site ensuring a single integration event per cell. They also carry a co-cistronic mCherry ORF downstream of the ORF, translated via an internal ribosome entry site. After integration, the ORF is expressed from the landing pad’s EF1ɑ promoter, and co-cistronic mCherry expression (and loss of BFP expression) allows for selection of integrants by FACS.

### Small-scale STK11 HeLa cell proliferation assay

The *STK11* variants tested in the small-scale assay were generated by site-directed mutagenesis. Briefly, forward and reverse primers harbouring a single base-pair mismatch were hybridized to wild-type *STK11* in pDONR223, amplified by PCR, then re-circularization of the plasmid by ligation completed a new *STK11-*variant pDONR223 plasmid. Wild-type and *STK11* variants were cloned into Bxb1-compatible destination plasmids and transfected as an equimolar pool into HeLa cells. A pool of successful *STK11*-integrant cells expressing the mCherry fluorescent marker were isolated by FACS and grown to confluency before collecting cells for the non-select control gDNA extraction. Over the following two weeks, HeLa cells were grown to confluency and harvested every three days for gDNA extraction. From each time-point, gDNA was prepared for Illumina short-read sequencing by amplifying the Bxb1 locus using forward and reverse primers designed to flank each of the *STK11* variants. Each variant tile at each time-point was sequenced at a depth of at least 10,000 reads, ensuring accurate measurements of each variant in the pooled population over time.

### Construction of codon randomised STK11 variant libraries

As a first step of the TileSeq framework, a modified version of pooled random-codon mutagenesis method (Precision Oligo-Pool based Code Alteration or POPCode)^65^ was used to construct the *STK11* variant libraries. Mutagenesis was separately applied in three regions of the codon-optimized *STK11* ORF, each encoding a ∼50 amino acid segment, following each step of the framework below to yield three distinct regionally-mutagenized libraries, each encoding full-length (1302 bp) *STK11*. Targeted mutagenesis was achieved with a pool of 28-38 bp-long oligonucleotides, each designed to have optimal melting temperatures using the POPcode oligo suite tool and each with a central NNN-degeneracy corresponding to a targeted codon. Collectively, each pool covered the length of the targeted *STK11* coding sequence region.

Oligonucleotides for each region were pooled and phosphorylated, and phosphorylated. Oligonucleotides and an upstream tagged primer were annealed to a codon-optimized *STK11* coding sequence (Supplementary Table 2) in the pDONR223 plasmid, by: 1) incubating the mix of oligonucleotides, primers, and plasmids at 95° for 3 minutes, then cooling down to 4°C by 0.1°C per second, to denature the plasmid and allow oligonucleotide and primer binding, followed by; 2) incubating with 2x Phusion Mastermix, 95° for 3 minutes cooling down to 4°C by 0.1°C per second, followed by 2 hours at 50°C to allow for oligonucleotide and primer elongation across the *STK11* sequence, then; 3) incubate with Taq DNA ligase at 45°C for 20 minutes to allow for ligation between oligonucleotide and primer extensions, creating new linear double-stranded DNA containing *STK11* variants, and; 4) incubate with primers that selectively target the upstream tag incorporated into new variant clones, removing unaltered wild-type *STK11* plasmids.

The newly POPCode-mutagenized strands were then amplified with primers containing flanking Gateway attB sites. The mutagenized attB-PCR products were next cloned en masse into the Entry vector pDONR223 by Gateway BP reactions, generating regional Entry libraries of ∼500,000 clones per region, with the goal of having each codon change represented by an average of 50 or more independent clones. Entry libraries were then transferred to a Gateway Destination vector (pDest_HC_rec_bxb_V2) compatible with the Bxb1 mammalian cells by en masse Gateway LR reactions, generating regional integration libraries from >500,000 transformants per region in order to maintain pool complexity. Both Gateway Entry and integration library transformations used NEB5α E. coli cells (NEB), with selection on LB agar plates using spectinomycin and ampicillin, respectively. Both Gateway Entry and Destination integration libraries were grown on 245 x 245mm^2^ bioassay dishes (Corning).

### Multiplexed assay for STK11 variant function

The HeLa-cell based proliferation assay used to evaluate *STK11* missense variants was performed in triplicate, with each of the three *STK11* regions transfected into cells three separate times (i.e. nine separate transfections in total). For each transfection, ∼15 million HeLa cells were transfected with a mix of Bxb1-integrase and the *STK11* variant library using Neon electroporation (1050V, 35ms, 2 pulses). We estimated ∼20-40% cell death and integration for ∼5-10% of remaining cells, yielding ∼500,000 independent integrant clones for each transfection of each region. Four to five days after transfection, cells were sorted by FACS to isolate the BFP-negative and mCherry-positive integrant cells that express STK11, then expanded in culture over the course of approximately two weeks. Cells were then collected for gDNA extraction and amplification using primers flanking the integrated *STK11* ORF.

### Quantifying variant abundance

Each region was conceptually divided into four tiles, each ∼110 nucleotides long to enable duplex sequencing (i.e., sequencing of both strands from each cluster on an Illumina flow cell) which in turn reduces base-calling error rates. Each tile was amplified from the *STK11* amplicon pool derived from each replicate pool of cells, under both non-selective and selective conditions, amplifying targeted tiles with primers carrying an Illumina sequencing-adaptor binding site. Next, a unique Illumina sequencing adapter was added to each tile via a subsequent “indexing PCR”. Equal amounts of the tiled indexed PCR products were pooled together and the resulting pooled library of an expected size of ∼300bp was purified using a 4%EX e-gel (Life Technologies) followed by MinElute Gel Extraction (Qiagen). After the library quantification via NEBNext Library Quant Kit (NEB), paired-end sequencing was performed on the tiles of each region with a sequencing depth of ∼2 million reads per tile using an Illumina NextSeq 500 instrument via a NextSeq 500/550 High Output Kit v2.

### Deriving functional scores for variants

An analysis pipeline, called TileseqMave (version 1.0.0), was used for the generation of the variant effect map (code, installation instructions and documentation are available on Github at https://github.com/rothlab/tileseqMave. TileseqMave uses bowtie2)^66^ to align both forward and reverse reads to the coding reference sequence. For each divergent base-call, the posterior probability of being a true variant was calculated based on the combination of base-call Q-scores. Variants with posterior probabilities above 90% were counted. For each codon change, a “marginal count” (number of times the change was observed irrespective of other co-occurring variants) was calculated. To derive the frequency of each codon change, the marginal count for each mutation was normalised by its “effective sequencing depth”, i.e. the number of reads deemed to have sufficient read quality on both strands to enable base-calling. Then, an error-corrected enrichment log-ratio was calculated by subtracting the frequency of the variant in the corresponding wild-type control from the post- and pre-selection frequencies and then calculating the log ratio post- to pre-selection frequencies. (Where multiple sequencing runs were required to obtain sufficient read counts, we subtracted counts from matched wild-type control libraries before aggregating counts.) Finally, the enrichment log-ratio was rescaled such that, after rescaling, synonymous variants have a median score of 1 and nonsense variants a median score of 0. For each variant, measurement error was regularised^67^ and propagated via bootstrapping.

To include only well-measured results, we filtered out variants with frequency below 2×10^−5^ and whose pre-selection frequencies were statistically indistinguishable from those in the wild-type control. We also removed variants for which sequencing replicates diverged by more than three times the amount expected based on the Poisson variance estimated from read count.

Further filtering to isolate high-quality variants was performed for the clinical analysis by removing variants where a variant’s confidence interval (score plus and minus estimated standard error) overlapped with the midpoint between synonymous and nonsense (i.e. a score of 0.5).

### Combining replicates to generate a unified functional score

For each biological replicate, after generating functional scores as described above for each codon substitution, we aggregated scores for codons corresponding to the same amino acid. Given that, both for Peutz-Jeghers and somatic cancer-causing *STK11* variants, pathogenic variants are thought to follow a loss-of-function mechanism, we sought to reduce the influence of outliers with scores greater than the median of synonymous variants (i.e., scores greater than 1), while still preserving rank-order of these scores. Therefore, scores above 1 were transformed using X’ = 1+[(X-1)*(1/X)] before averaging. We next considered whether measurements for different codons agreed with respect to whether the variant was damaging (<0.5 score) or tolerated (>0.5 score). For substitutions with two codons measured, we filtered out amino acid substitutions that disagreed on damaging vs tolerated, if the difference in scores was >0.3. For substitutions with three or more codons measured, the median score was taken across all codons.

Next, we sought to integrate the codon-merged scores across the three biological replicates (i.e. independent transfections). The same process as above for assessing agreement and combining replicate scores was implemented, except that variants without biological replicate measurements, i.e., with a well-measured score from only one transfection, were removed.

### Conservation, solvent accessibility, and protein stability

To quantify evolutionary conservation, we used ConSurf^29^ to assign a conservation score to each position along the protein. To enable ConSurf’s use of structural homology when choosing homologs for comparison, we provided a predicted structure [using AlphaFold version 2022-11-01, Monomer v2.0 pipeline]^31^. The median functional score for every conserved position (defined by ConSurf as having a score <-.47) was compared to functional scores in non-conserved positions (defined by ConSurf as having score >.91) using a Wilcoxon rank-sum test.

Solvent accessibility was calculated using FreeSASA [version 2.02 October 22nd, 2017] using the above-mentioned pdb structure^30^. Positions with relative side-chain solvent accessibility <20% were considered inaccessible and those with >50% were considered accessible. ΔΔG thermostability predictions were made for all possible missense variants using the FoldX software^28^, such that ΔΔG scores above 0 are more likely to destabilise the protein while scores below 0 are more likely to retain structure.

### Reference set of clinically annotated variants

A set of clinically-annotated variants was provided via the Ambry Genetics variant classification system which captures evidence of strength in terms of points towards pathogenicity or benignity^9^. Variants with at least six pathogenic points were included in the positive reference set (including variants annotated as either pathogenic and likely pathogenic variants) and variants with at least two benign points were included in the negative reference set (including variants annotated as either benign or likely benign).

### Calibrating log-likelihood ratios to ACMG evidence strengths

To derive estimates of the strength of evidence towards pathogenicity annotation that are both compatible with a Bayesian clinical variant classification approach and are continuously quantitative unlike points-based approaches, we estimated the log-likelihood ratio of pathogenicity (LLR) for each variant as previously described^39^. Briefly, probability density functions were separately estimated for scores from the positive and negative reference variant sets using kernel density estimation (using a Gaussian kernel with a bandwidth determined by biased cross-validation). Then, for each variant, the LLR could be calculated as the log of the ratio of the two probability densities, but ‘regularised’ towards the uniform distribution to avoid extreme LLRs in score ranges with fewer reference variants. LLR values were then compared to threshold values defined by the ACMG/AMP variant classification framework using the strategy of Tavtigian et al.^40^ as subsequently adapted^39^.

## Supporting information

All_STK11_Functional_Scores

## Acknowledgements

The authors gratefully acknowledge funding from the National Human Genome Research Institute of the National Institutes of Health (NIH/NHGRI) Center of Excellence in Genomic Science (CEGS) Initiative (HG010461), the NIH/NHGRI Impact of Genomic Variation on Function (IGVF) Initiative (HG011989), and the Canadian Institutes of Health Research (FDN-159926).

## Data Availability

All *STK11* variant functional scores generated in this study are provided in the supplemental csv file named ‘All_*STK11*_Functional_Scores’ and are also available online at MaveDB (https://www.mavedb.org), accession number: urn:mavedb:00001246-a-1.

## Supplementary Figures

**Supplementary Figure 1.**
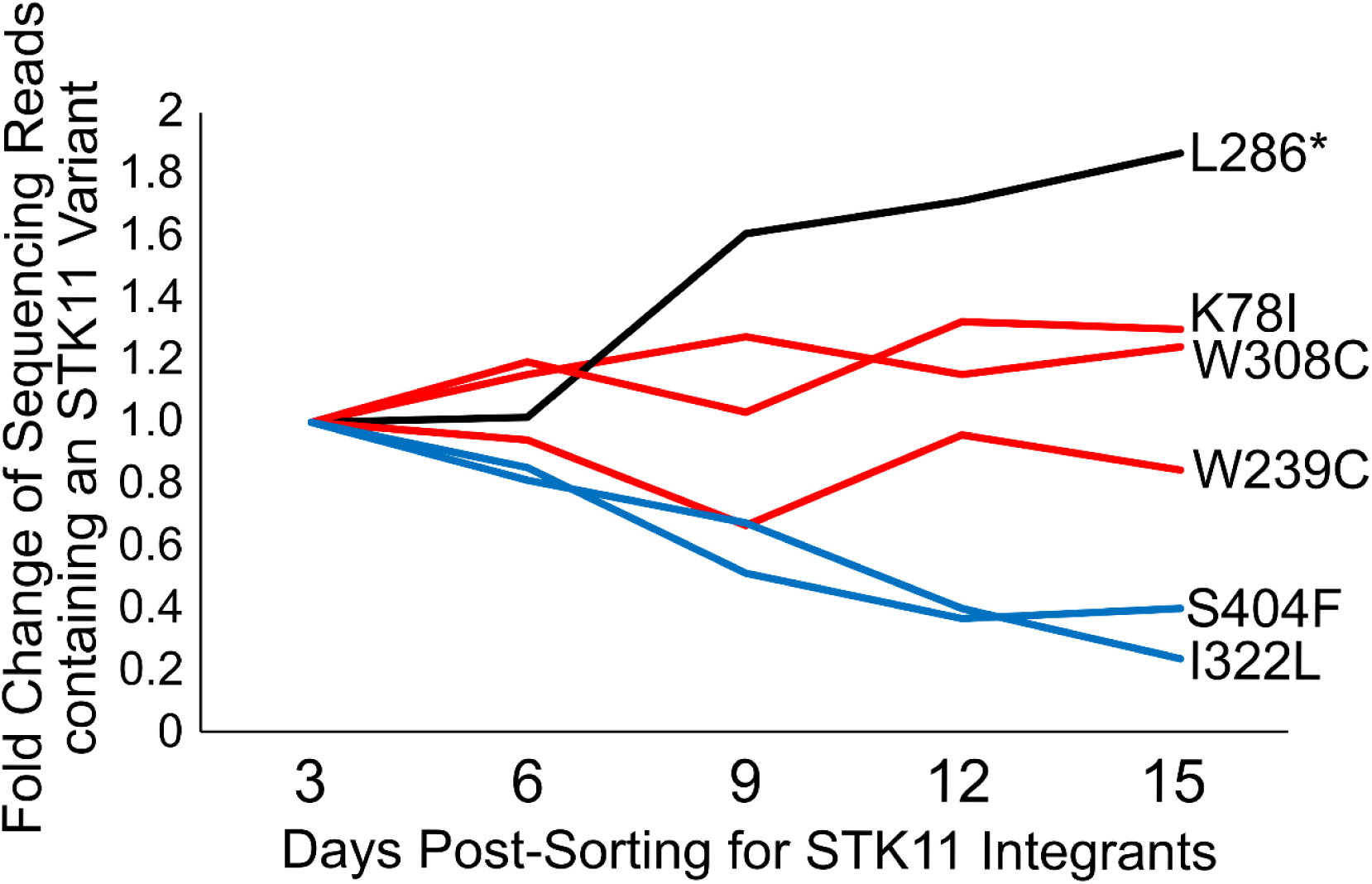
Small-Scale Validation of the HeLa Proliferation Assay. Equimolar quantities of Bxb1 plasmids bearing *STK11* variants (K78I, W239C, L286*, W308C, I322L, and S404F) were transfected into HeLa cells for stable integration. Successful *STK11* integrants were isolated by FACS then grown in culture over two-week time with a portion of the cells harvested every three days for sequencing. As a proxy for proliferation rate, the change in frequency of each *STK11* variant in the pool was quantified by NGS at multiple time-points. Missense variants expected to impact protein function are shown in red, nonsense variants in black, and tolerated missense in blue.

**Supplementary Figure 2.**
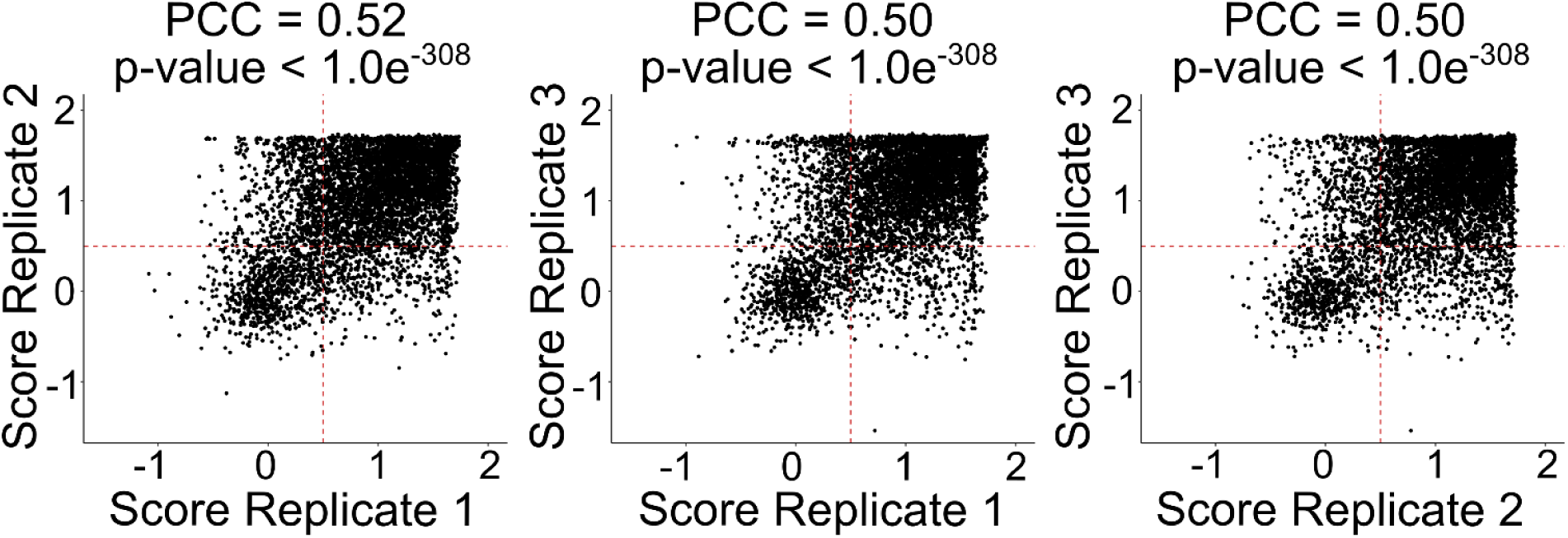
Correlation of Functional Scores Across Biological Replicates. Missense variant functional scores from each of the three biological replicates (i.e. independent transfections) were plotted in pairwise combinations. Intermediate functional scores of 0.5 are indicated by horizontal and vertical red dashed lines. Pearson correlation was calculated, and the correlation coefficient and p-value are reported above the plots.

**Supplementary Figure 3.**
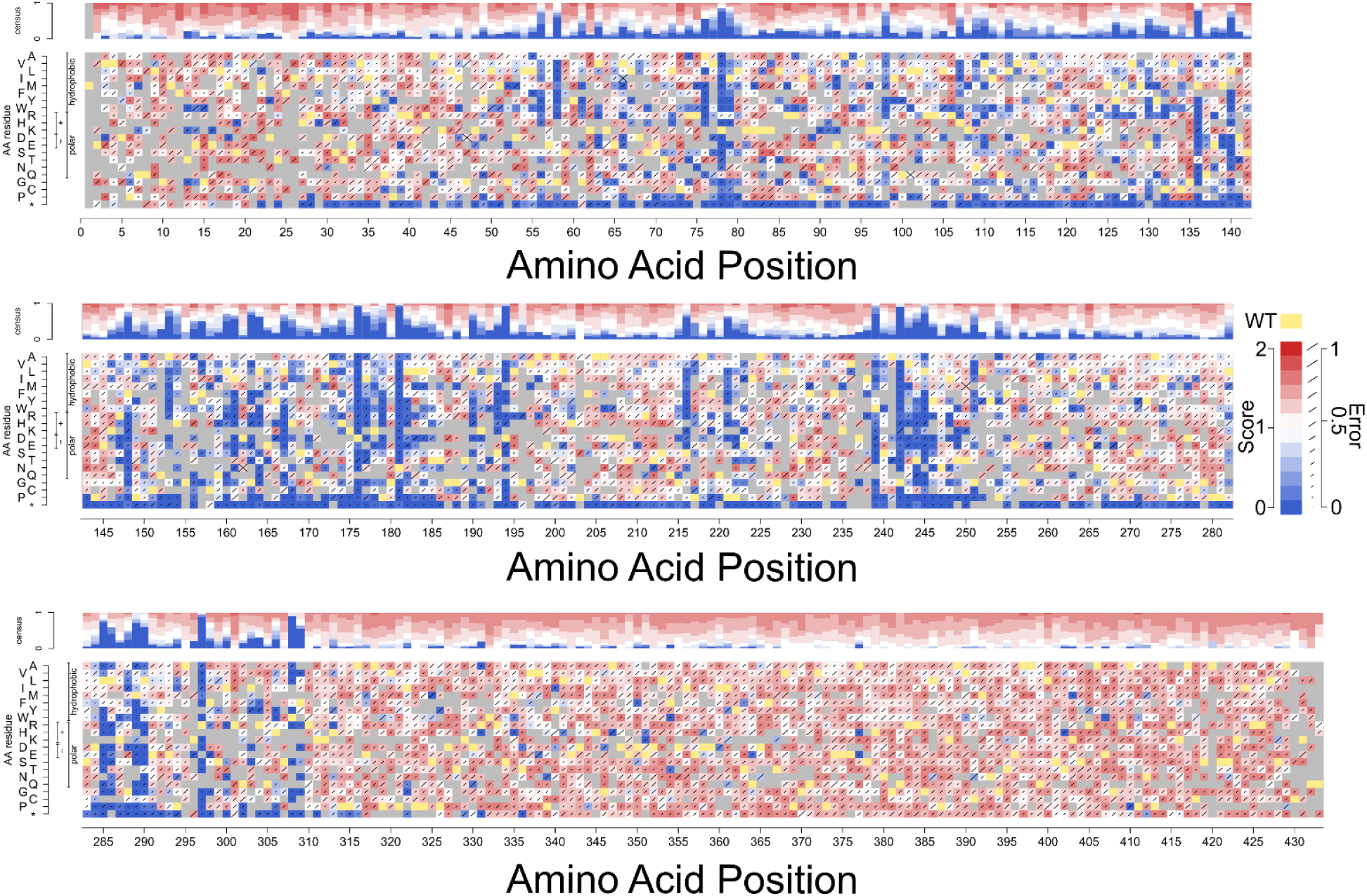
Full-Length STK11 Variant Effect Map. Functional scores for all well-measured variants are shown as a heatmap with all possible amino acid substitutions on the y-axis, and position along STK11 on the x-axis. Yellow indicates the wild-type residue, gray for variants that did not pass filtering, blue for damaging scores near 0, white for tolerated scores near 1, red for scores above 1. Error rates (calculated as described in the methods) are shown as diagonal lines within each cell, with larger lines for higher error rates.

**Supplementary Figure 4.**
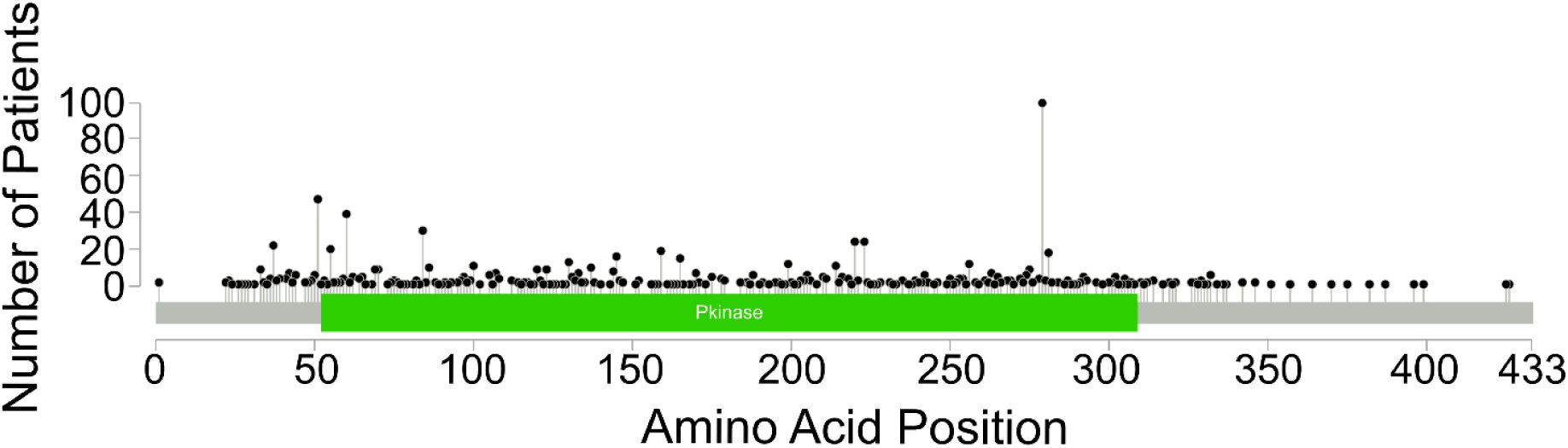
STK11 Truncating Variants on cBioPortal. The curated set of non-redundant studies was selected on cBioPortal (As of May 2025) with a gene-specific query for *STK11*. All unique *STK11* truncating variants were depicted as black dots on the lollipop plot, with the x-axis showing position along STK11, and the y-axis showing the number of patients with each variant.

**Supplementary Figure 5.**
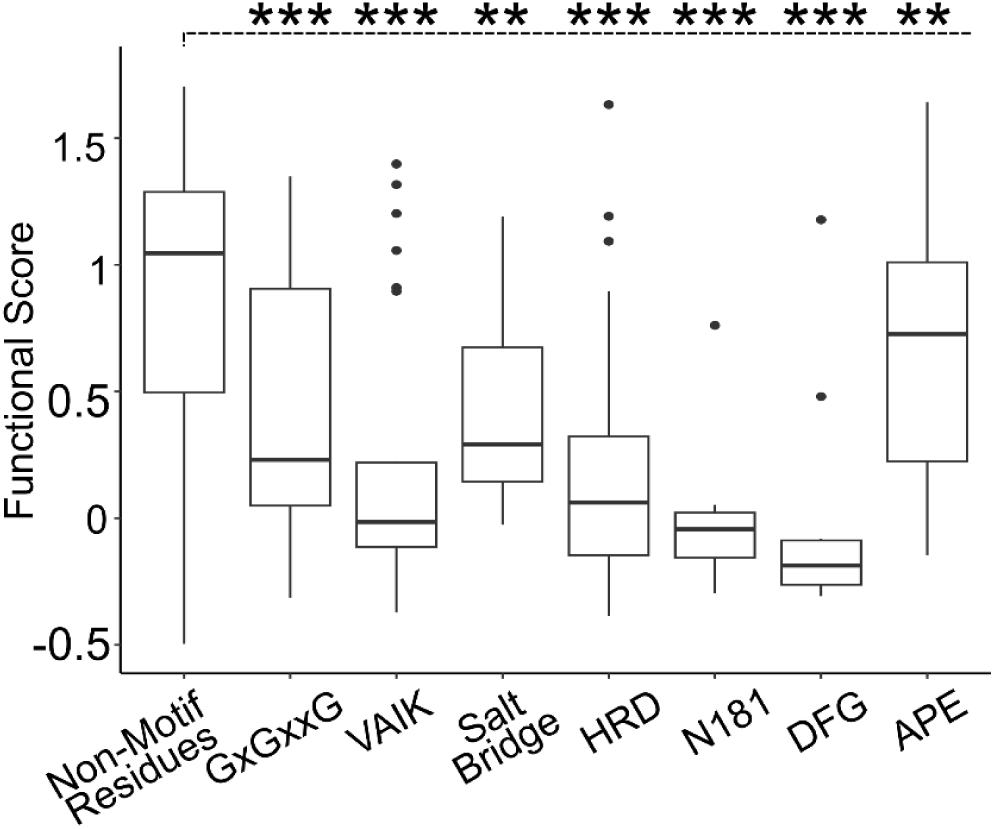
Functional Scores at Conserved Motif Positions. Functional score for each variant at conserved kinase motif positions that matched the consensus sequence are shown. Boxes show the interquartile range between the 25th to 75th quartiles, with a solid horizontal line for the 50th percentile. Vertical lines extending from each box represent variants with scores within 1.5 times the interquartile range, and circular dots for variants exceeding that threshold.

**Supplementary Figure 6.**
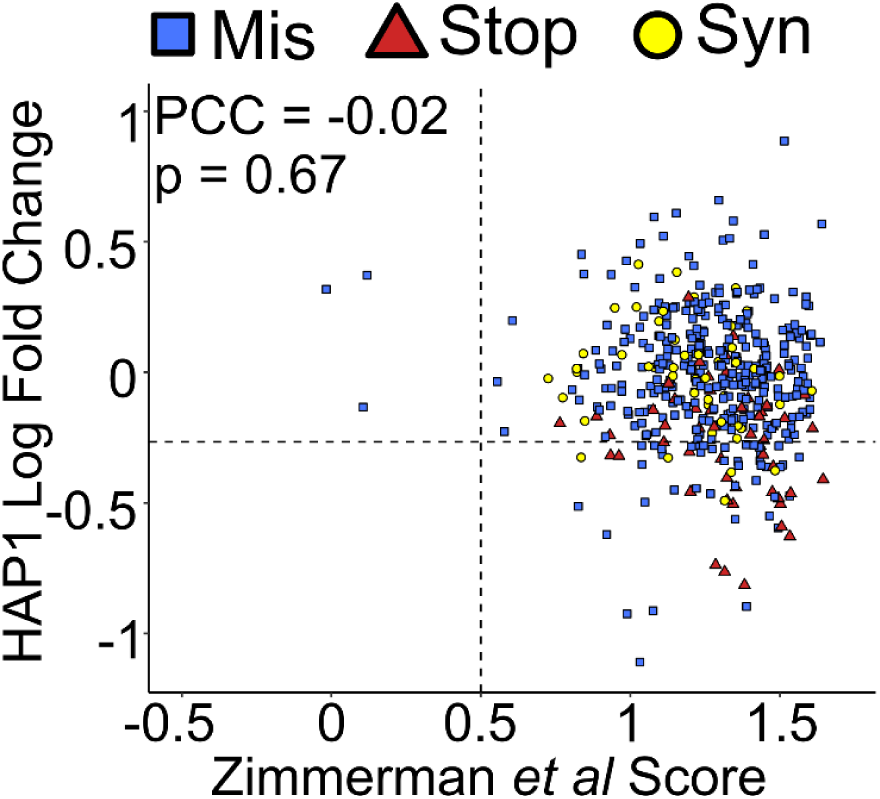
*STK11* Exon 9 Essentiality Assay in HAP1 Cells. Functional scores in exon 9 were matched to results from a *STK11* essentiality assay in HAP1 cells. Loss of *STK11* function in HAP1 cells reduces their ability to proliferate and survive, leading to drop-out in the pooled population. Missense variants are shown as blue squares, nonsense variants as red triangles, and synonymous variants as yellow circles.

**Supplementary Figure 7.**
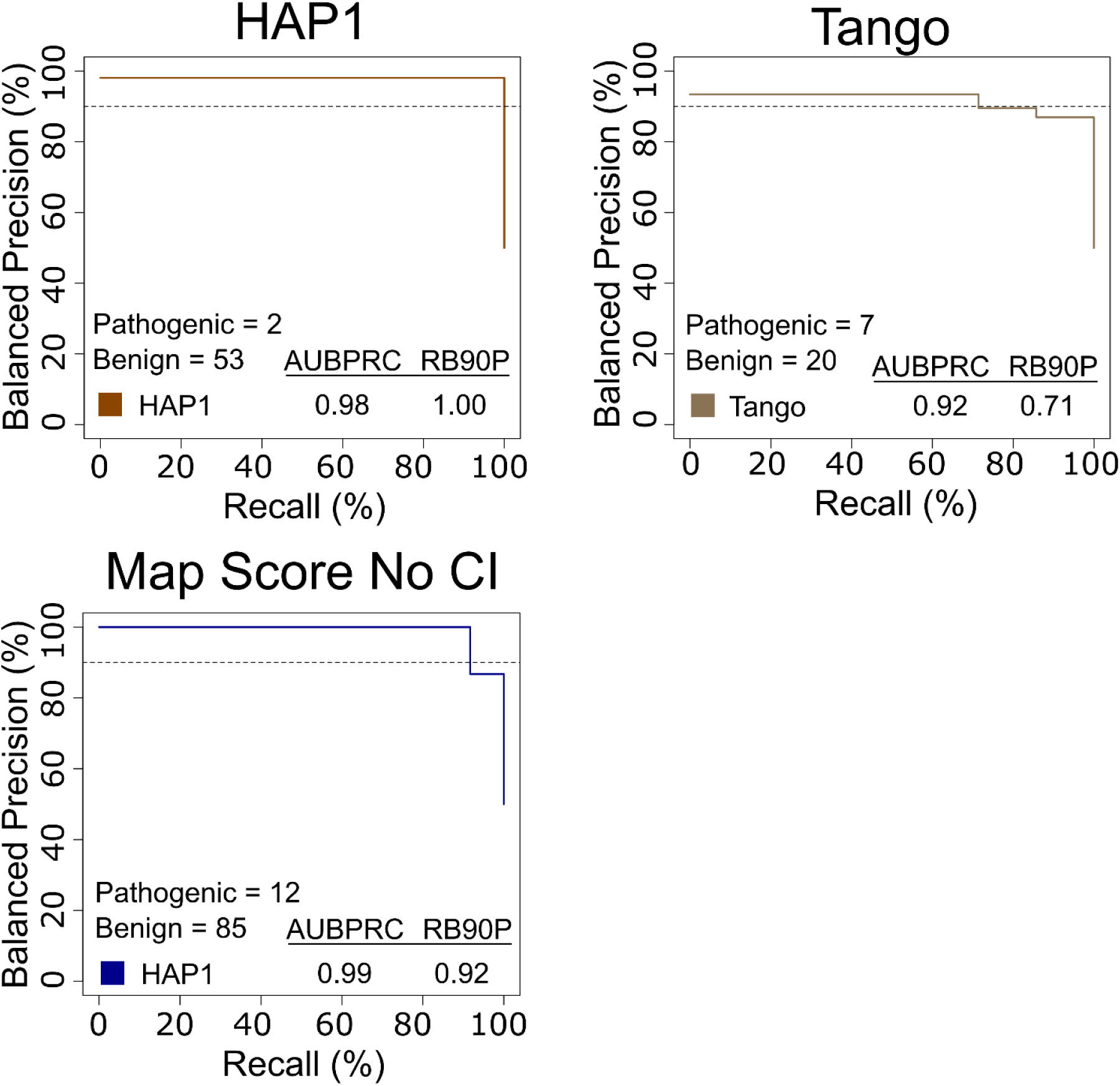
Precision-Recall Curves for Alternative Assays. Precision-recall curves were generated for *STK11* functional scores from the HAP1 essentiality assay, the Tango xenograft, and all HeLa map scores without the confidence interval filter. Using a known set of clinically annotated pathogenic or benign *STK11* variants from Ambry Genetics, we evaluated balanced precision—defined at each score threshold by the fraction of variants that are pathogenic given a balanced (50% prior probability of pathogenicity) test set—versus recall (fraction of pathogenic variants captured at this threshold). The horizontal dashed line indicates R90BP with the numerical AUPRC and R90BP listed in the bottom-left hand legend.

**Supplementary Figure 8.**
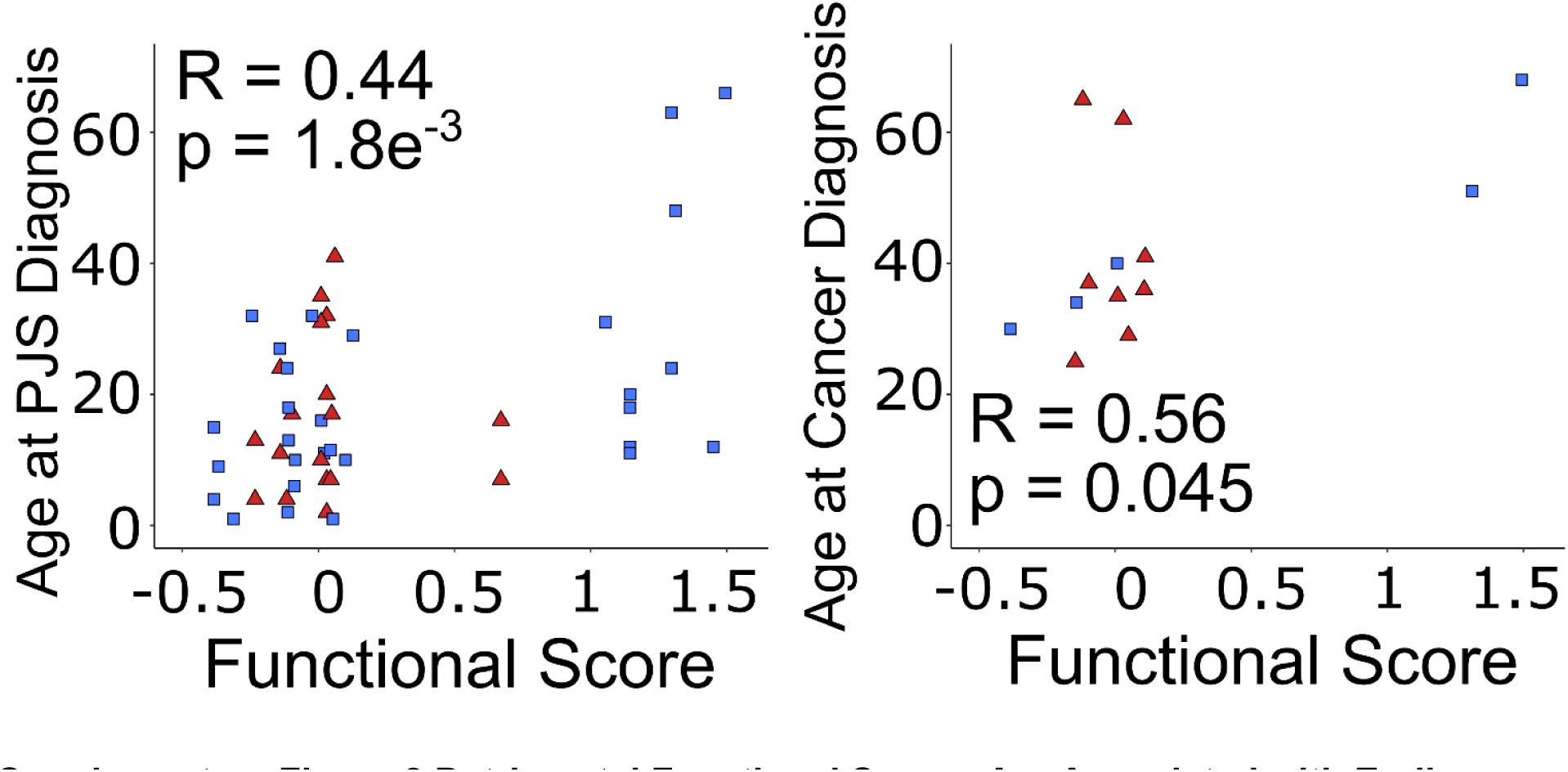
Detrimental Functional Scores Are Associated with Earlier Diagnosis of PJS and Onset of Cancer. Age at initial PJS diagnosis and age at PJS-associated cancer were collected from patient charts at the Zane Cohen Centre as well as from a literature review of available PJS case studies. Red triangles indicate nonsense variants, blue squares represent missense variants. The age at first diagnosis was determined by the earliest manifestation of recorded symptoms (such as hyper-pigmentation, observation of intestinal polyps, intussusception, etc.). Pearson correlation between age at PJS diagnosis or age at PJS-associated cancer and functional score is provided.

**Supplementary Figure 9.**
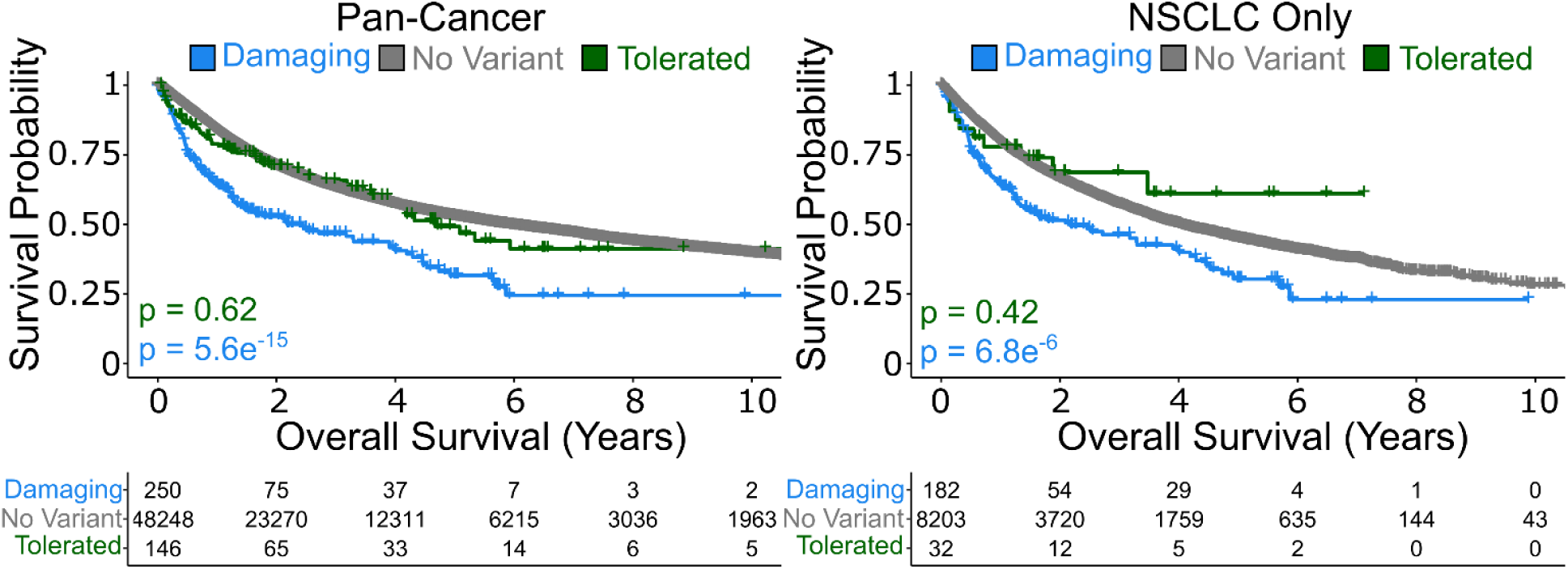
Stratifying Somatic Cancers by Functional Score in cBioPortal. Metrics relating to overall survival for individuals with cancer were extracted from a non-redundant set of clinical studies on cBioPortal (as of March 2025). Datasets on cBioPortal directly provided metrics of overall survival, here displayed in years. Patients were stratified into those without an *STK11* variant (gray), a tolerated *STK11* variant (green), or a damaging *STK11* variant (blue) according to our map score. Kaplan-Meier survival curves visualized the probability of survival over time in years, each ‘drop’ representing a death end-point, and each ‘tick’ a currently living individual. Log-rank tests were performed to compare outcomes for individuals with a damaging or tolerated *STK11* variant against those with wild-type *STK11*.

## Supplementary Tables

**Supplementary Table 1.**
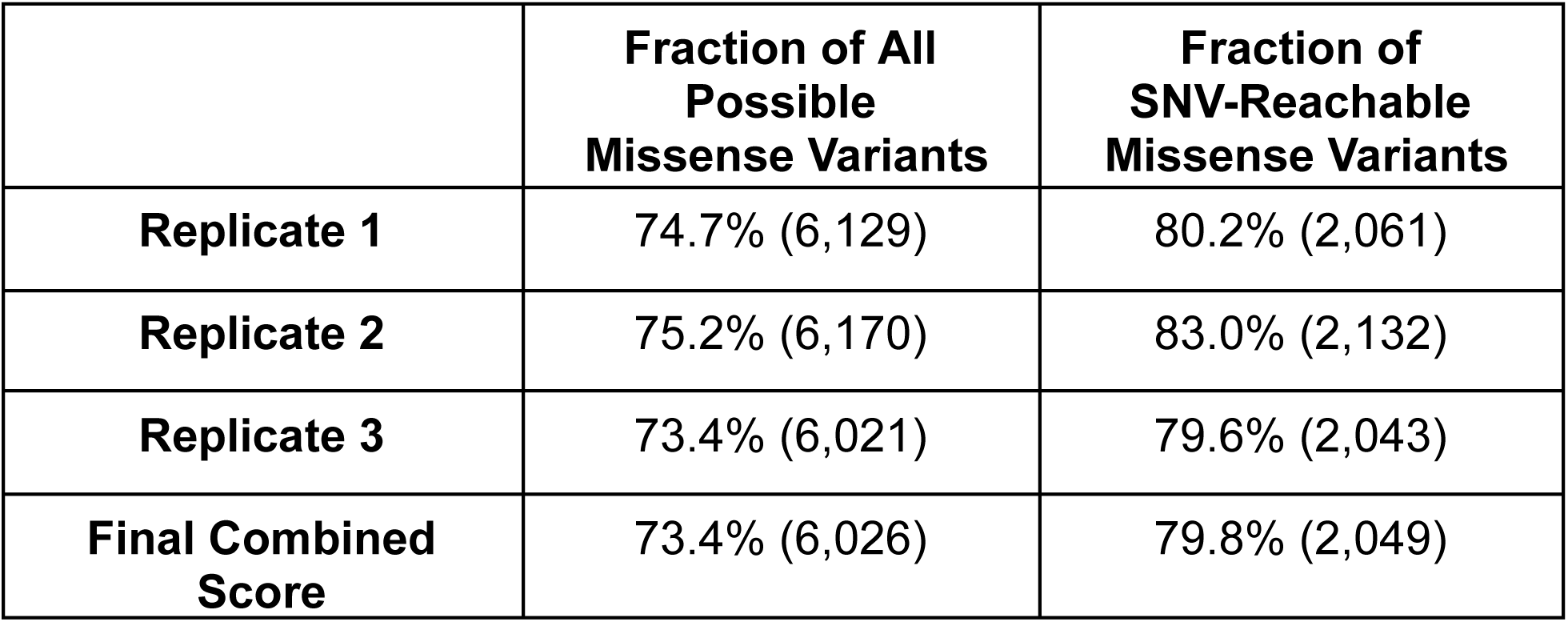
Coverage of *STK11* Missense Variants in Each Biological Replicate. Each biological replicate consists of three independent transfections (one per mutagenesis region of *STK11*) of the HeLa proliferation assay. The fraction of all possible *STK11* missense variants (8,208 possible variants), and the fraction of single-nucleotide accessible variants (2,568 possible variants) is shown along with the number of variants in brackets.

**Supplementary Table 2.**
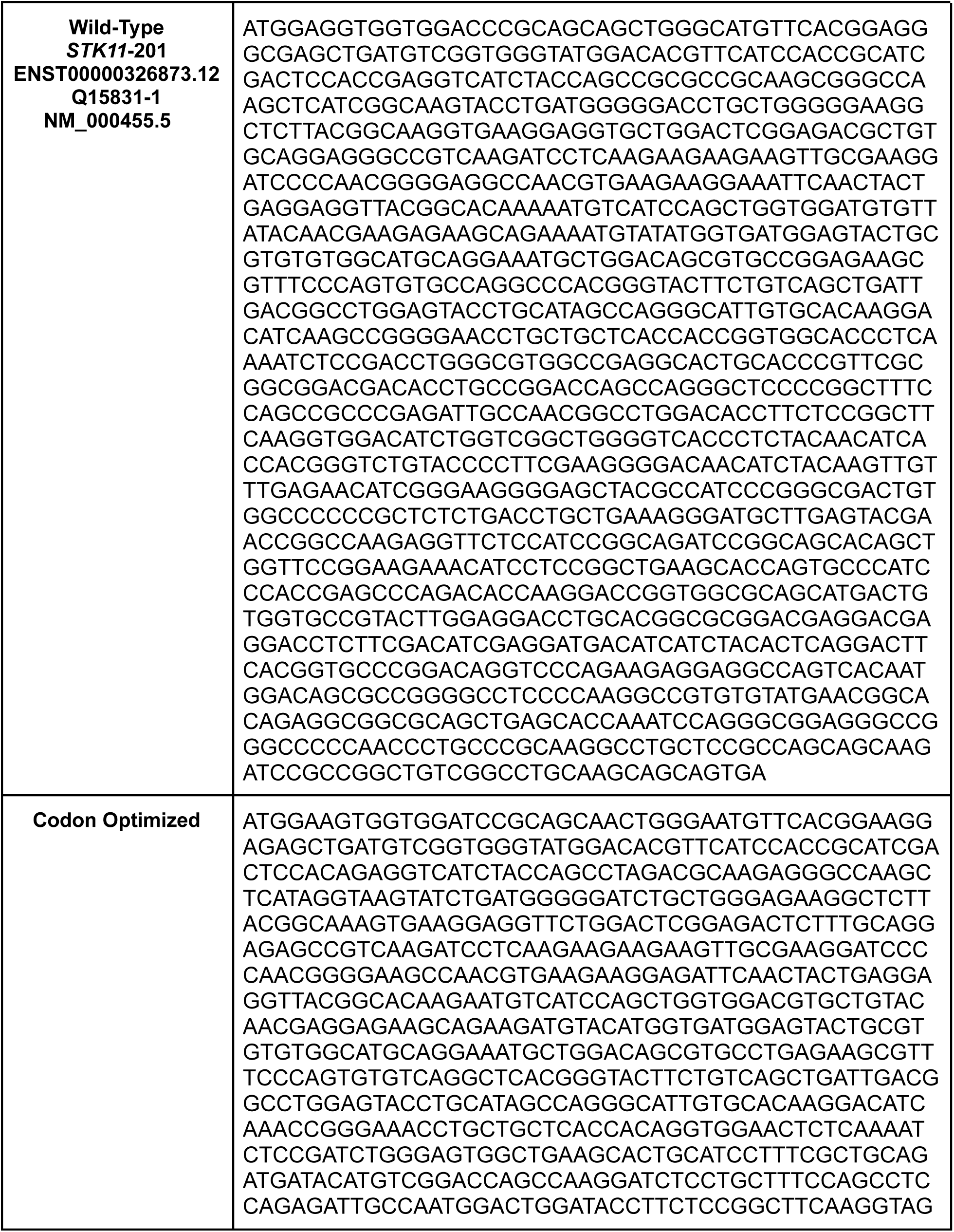
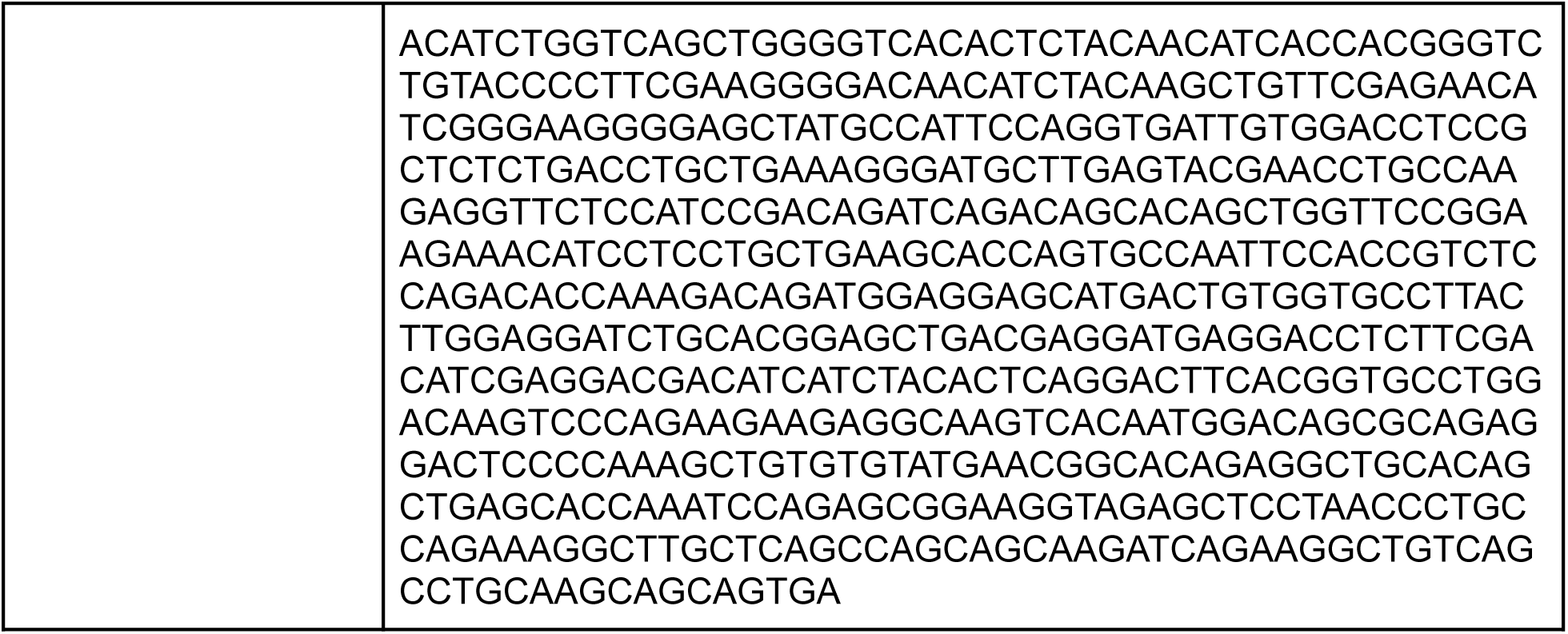
Codon Optimized *STK11* cDNA Sequence. To facilitate mutagenesis of *STK11* (Transcript ENST00000326873.12, RefSeq NM_000455.5), we codon-optimized to ensure GC-content was consistently between 40-60% for each 33bp stretch across the gene.

## References

1. Bourouh, M. & Marignani, P. A. The Tumor Suppressor Kinase LKB1: Metabolic Nexus. Front. Cell Dev. Biol. 10, 881297 (2022).

2. Pons-Tostivint, E., Lugat, A., Fontenau, J.-F., Denis, M. G. & Bennouna, J. STK11/LKB1 Modulation of the Immune Response in Lung Cancer: From Biology to Therapeutic Impact. Cells 10, 3129 (2021).

3. Skoulidis, F. et al. STK11/LKB1 Mutations and PD-1 Inhibitor Resistance in KRAS-Mutant Lung Adenocarcinoma. Cancer Discov. 8, 822–835 (2018).

4. Giardiello, F. M. & Trimbath, J. D. Peutz-Jeghers Syndrome and Management Recommendations. Clin. Gastroenterol. Hepatol. 4, 408–415 (2006).

5. Jelsig, A. M. et al. Survival, surveillance, and genetics in patients with Peutz-Jeghers syndrome: A nationwide study. Clin. Genet. 104, 81–89 (2023).

6. Beggs, A. D. et al. Peutz–Jeghers syndrome: a systematic review and recommendations for management. Gut 59, 975–986 (2010).

7. Landrum, M. J. et al. ClinVar: improving access to variant interpretations and supporting evidence. Nucleic Acids Res. 46, D1062–D1067 (2018).

8. Richards, S. et al. Standards and guidelines for the interpretation of sequence variants: a joint consensus recommendation of the American College of Medical Genetics and Genomics and the Association for Molecular Pathology. Genet. Med. 17, 405–424 (2015).

9. Fayer, S. et al. Closing the gap: Systematic integration of multiplexed functional data resolves variants of uncertain significance in BRCA1, TP53, and PTEN. Am. J. Hum. Genet. 108, 2248–2258 (2021).

10. Scott, A. et al. Saturation-scale functional evidence supports clinical variant interpretation in Lynch syndrome. Genome Biol. 23, 266 (2022).

11. Shirts, B. H., Pritchard, C. C. & Walsh, T. Family-Specific Variants and the Limits of Human Genetics. Trends Mol. Med. 22, 925–934 (2016).

12. Findlay, G. M. et al. Accurate classification of BRCA1 variants with saturation genome editing. Nature 562, 217–222 (2018).

13. Buckley, M. et al. Saturation genome editing maps the functional spectrum of pathogenic VHL alleles. Nat. Genet. 56, 1446–1455 (2024).

14. McCabe, M. T., Powell, D. R., Zhou, W. & Vertino, P. M. Homozygous deletion of the STK11/LKB1 locus and the generation of novel fusion transcripts in cervical cancer cells. Cancer Genet. Cytogenet. 197, 130–141 (2010).

15. Mack, H. I. D. & Munger, K. The LKB1 tumor suppressor differentially affects anchorage independent growth of HPV positive cervical cancer cell lines. Virology 446, 9–16 (2013).

16. Nguyen, H. B., Babcock, J. T., Wells, C. D. & Quilliam, L. A. LKB1 tumor suppressor regulates AMP kinase/mTOR-independent cell growth and proliferation via the phosphorylation of Yap. Oncogene 32, 4100–4109 (2013).

17. Zhang, X., Chen, H., Wang, X., Zhao, W. & Chen, J. J. Expression and transcriptional profiling of the LKB1 tumor suppressor in cervical cancer cells. Gynecol. Oncol. 134, 372–378 (2014).

18. Granado-Martínez, P. et al. STK11 (LKB1) missense somatic mutant isoforms promote tumor growth, motility and inflammation. *Commun*. Biol. 3, 1–14 (2020).

19. Matreyek, K. A., Stephany, J. J., Chiasson, M. A., Hasle, N. & Fowler, D. M. An improved platform for functional assessment of large protein libraries in mammalian cells. Nucleic Acids Res. 48, e1 (2020).

20. Mehenni, H. et al. Loss of LKB1 kinase activity in Peutz-Jeghers syndrome, and evidence for allelic and locus heterogeneity. Am. J. Hum. Genet. 63, 1641–1650 (1998).

21. Islam, Md. J., Khan, A. M., Parves, Md. R., Hossain, M. N. & Halim, M. A. Prediction of Deleterious Non-synonymous SNPs of Human STK11 Gene by Combining Algorithms, Molecular Docking, and Molecular Dynamics Simulation. Sci. Rep. 9, 16426 (2019).

22. Hartley, J. L., Temple, G. F. & Brasch, M. A. DNA Cloning Using In Vitro Site-Specific Recombination. Genome Res. 10, 1788–1795 (2000).

23. Kochetov, A. V. Alternative translation start sites and hidden coding potential of eukaryotic mRNAs. BioEssays News Rev. Mol. Cell. Dev. Biol. 30, 683–691 (2008).

24. Zeqiraj, E., Filippi, B. M., Deak, M., Alessi, D. R. & van Aalten, D. M. F. Structure of the LKB1-STRAD-MO25 complex reveals an allosteric mechanism of kinase activation. Science 326, 1707–1711 (2009).

25. Hudson, A. M. et al. Truncation- and motif-based pan-cancer analysis reveals tumor-suppressing kinases. Sci. Signal. 11, eaan6776 (2018).

26. Lahiry, P., Torkamani, A., Schork, N. J. & Hegele, R. A. Kinase mutations in human disease: interpreting genotype–phenotype relationships. Nat. Rev. Genet. 11, 60–74 (2010).

27. Varga, M. J., Richardson, M. E. & Chamberlin, A. Structural biology in variant interpretation: Perspectives and practices from two studies. Am. J. Hum. Genet. 112, 984–992 (2025).

28. Schymkowitz, J. et al. The FoldX web server: an online force field. Nucleic Acids Res. 33, W382–W388 (2005).

29. Ben Chorin, A., et al. ConSurf-DB: An accessible repository for the evolutionary conservation patterns of the majority of PDB proteins. Protein Sci. Publ. Protein Soc. 29, 258–267 (2020).

30. Mitternacht, S. FreeSASA: An open source C library for solvent accessible surface area calculations. F1000Research 5, 189 (2016).

31. Jumper, J. et al. Highly accurate protein structure prediction with AlphaFold. Nature 596, 583–589 (2021).

32. Boudeau, J. et al. Analysis of the LKB1-STRAD-MO25 complex. J. Cell Sci. 117, 6365–6375 (2004).

33. Boudeau, J. et al. Functional analysis of LKB1/STK11 mutants and two aberrant isoforms found in Peutz-Jeghers Syndrome patients. Hum. Mutat. 21, 172–172 (2003).

34. Chen, L. et al. A sensitive NanoString-based assay to score STK11 (LKB1) pathway disruption in lung adenocarcinoma. J. Thorac. Oncol. Off. Publ. Int. Assoc. Study Lung Cancer 11, 838–849 (2016).

35. Lacoste, J. et al. Pervasive mislocalization of pathogenic coding variants underlying human disorders. Cell 187, 6725–6741.e13 (2024).

36. Obolenski, S., Olvera-León, R., Sun, D., Adams, D. J. & Waters, A. J. Protocol for the functional evaluation of genetic variants using saturation genome editing. STAR Protoc. 6, 103710 (2025).

37. Baas, A. F. et al. Activation of the tumour suppressor kinase LKB1 by the STE20-like pseudokinase STRAD. EMBO J. 22, 3062–3072 (2003).

38. Donnelly, L. L. et al. Functional assessment of somatic STK11 variants identified in primary human non-small cell lung cancers. Carcinogenesis 42, 1428–1438 (2021).

39. van Loggerenberg, W. et al. Systematically testing human HMBS missense variants to reveal mechanism and pathogenic variation. Am. J. Hum. Genet. 110, 1769–1786 (2023).

40. Tavtigian, S. V. et al. Modeling the ACMG/AMP Variant Classification Guidelines as a Bayesian Classification Framework. Genet. Med. Off. J. Am. Coll. Med. Genet. 20, 1054–1060 (2018).

41. Jiang, Y.-L. et al. The altered activity of P53 signaling pathway by STK11 gene mutations and its cancer phenotype in Peutz-Jeghers syndrome. BMC Med. Genet. 19, 141 (2018).

42. Resta, N. et al. Cancer risk associated with *STK11/LKB1* germline mutations in Peutz–Jeghers syndrome patients: Results of an Italian multicenter study. Dig. Liver Dis. 45, 606–611 (2013).

43. Amos, C. I. et al. Genotype-phenotype correlations in Peutz-Jeghers syndrome. J. Med. Genet. 41, 327–333 (2004).

44. Salloch, H. et al. Truncating mutations in Peutz-Jeghers syndrome are associated with more polyps, surgical interventions and cancers. Int. J. Colorectal Dis. 25, 97–107 (2010).

45. Chiang, J.-M. & Chen, T.-C. Clinical manifestations and STK11 germline mutations in Taiwanese patients with Peutz-Jeghers syndrome. Asian J. Surg. 41, 480–485 (2018).

46. Gu, G.-L. et al. Detection and analysis of common pathogenic germline mutations in Peutz-Jeghers syndrome. World J. Gastroenterol. 27, 6631–6646 (2021).

47. Lim, W. et al. Further observations on LKB1/STK11 status and cancer risk in Peutz–Jeghers syndrome. Br. J. Cancer 89, 308–313 (2003).

48. Mehenni, H. et al. Molecular and clinical characteristics in 46 families affected with Peutz-Jeghers syndrome. Dig. Dis. Sci. 52, 1924–1933 (2007).

49. Schumacher, V. et al. STK11 genotyping and cancer risk in Peutz-Jeghers syndrome. J. Med. Genet. 42, 428–435 (2005).

50. Zhao, H.-M. et al. Clinical and Genetic Study of Children With Peutz-Jeghers Syndrome Identifies a High Frequency of STK11 De Novo Mutation. J. Pediatr. Gastroenterol. Nutr. 68, 199–206 (2019).

51. Zheng, B. et al. A Clinical and Molecular Genetic Study in 11 Chinese Children With Peutz-Jeghers Syndrome. J. Pediatr. Gastroenterol. Nutr. 64, 559–564 (2017).

52. Cerami, E. et al. The cBio cancer genomics portal: an open platform for exploring multidimensional cancer genomics data. Cancer Discov. 2, 401–404 (2012).

53. AACR Project GENIE Consortium. AACR Project GENIE: Powering Precision Medicine through an International Consortium. Cancer Discov. 7, 818–831 (2017).

54. Wagner, A. et al. The Management of Peutz-Jeghers Syndrome: European Hereditary Tumour Group (EHTG) Guideline. J. Clin. Med. 10, 473 (2021).

55. Shire, N. J. et al. STK11 (LKB1) mutations in metastatic NSCLC: Prognostic value in the real world. PLOS ONE 15, e0238358 (2020).

56. Sanders, M. J. et al. Defining the mechanism of activation of AMP-activated protein kinase by the small molecule A-769662, a member of the thienopyridone family. J. Biol. Chem. 282, 32539–32548 (2007).

57. Devarakonda, S. et al. A phase II study of everolimus in patients with advanced solid malignancies with TSC1, TSC2, NF1, NF2 or STK11 mutations. J. Thorac. Dis. 13, 4054–4062 (2021).

58. Galan-Cobo, A. et al. LKB1 and KEAP1/NRF2 Pathways Cooperatively Promote Metabolic Reprogramming with Enhanced Glutamine Dependence in KRAS-Mutant Lung Adenocarcinoma. Cancer Res. 79, 3251–3267 (2019).

59. Caiola, E. et al. LKB1 Deficiency Renders NSCLC Cells Sensitive to ERK Inhibitors. J. Thorac. Oncol. Off. Publ. Int. Assoc. Study Lung Cancer 15, 360–370 (2020).

60. Patel, A. et al. Abstract 3916: TNG260, a small molecule CoREST inhibitor, sensitizes STK11-mutant NSCLC to anti-PD1 immunotherapy. Cancer Res. 84, 3916 (2024).

61. Li, H. et al. AXL targeting restores PD-1 blockade sensitivity of STK11/LKB1 mutant NSCLC through expansion of TCF1+ CD8 T cells. Cell Rep. Med. 3, 100554 (2022).

62. Bai, X. et al. CDK4/6 inhibition triggers ICAM1-driven immune response and sensitizes LKB1 mutant lung cancer to immunotherapy. Nat. Commun. 14, 1247 (2023).

63. Pore, N. et al. Resistance to Durvalumab and Durvalumab plus Tremelimumab Is Associated with Functional STK11 Mutations in Patients with Non-Small Cell Lung Cancer and Is Reversed by STAT3 Knockdown. Cancer Discov. 11, 2828–2845 (2021).

64. Yang, X. et al. A public genome-scale lentiviral expression library of human ORFs. Nat. Methods 8, 659–661 (2011).

65. Weile, J. et al. A framework for exhaustively mapping functional missense variants. Mol. Syst. Biol. 13, 957 (2017).

66. Langmead, B. & Salzberg, S. L. Fast gapped-read alignment with Bowtie 2. Nat. Methods 9, 357–359 (2012).

67. Baldi, P. & Long, A. D. A Bayesian framework for the analysis of microarray expression data: regularized t -test and statistical inferences of gene changes. Bioinforma. Oxf. Engl. 17, 509–519 (2001).

